# Target Recognition in Tandem WW Domains: Complex Structures for Parallel and Antiparallel Ligand Orientation in h-FBP21 Tandem WW

**DOI:** 10.1101/2021.11.22.469489

**Authors:** Marius T. Wenz, Miriam Bertazzon, Jana Sticht, Stevan Aleksić, Daniela Gjorgjevikj, Christian Freund, Bettina G. Keller

**Affiliations:** Institute for Chemistry and Biochemistry, Molecular Dynamics Group, Arnimallee 22, Freie Universität Berlin, 14195 Berlin, Germany; Institute for Chemistry and Biochemistry, Protein Biochemistry Group, Thielallee 63, Freie Universität Berlin, 14195 Berlin, Germany; Core Facility BioSupraMol, Takustr. 3, Freie Universität Berlin, 14195, Berlin, Germany; Medicinal Chemistry, Boehringer Ingelheim Pharma GmbH & Co. KG, 88397 Biberach, Germany

## Abstract

Protein-protein interactions often rely on specialized recognition domains, such as WW domains, which bind to specific proline-rich sequences. The specificity of these protein-protein interactions can be increased by tandem repeats, i.e. two WW domains connected by a linker. With a flexible linker, the WW domains can move freely with respect to each other. Additionally, the tandem WW domains can bind in two different orientations to their target sequences. This makes the elucidation of complex structures of tandem WW domains extremely challenging. Here, we identify and characterize two complex structures of the tandem WW domain of human formin-binding protein 21 and a peptide sequence from its natural binding partner, the core-splicing protein SmB/B’. The two structures differ in the ligand orientation, and consequently also in the relative orientation of the two WW domains. We analyze and probe the interactions in the complexes by molecular simulations and NMR experiments. The workflow to identify the complex structures uses molecular simulations, density-based clustering and peptide docking. It is designed to systematically generate possible complex structures for repeats of recognition domains. These structures will help us to understand the synergistic and multivalency effects that generate the astonishing versatility and specificity of protein-protein interactions.

## 1 Introduction

Molecular machines carry out the processes that constitute life.^1,2^ These machines are multiprotein or multi-protein-RNA complexes,^3–5^ such as the spliceosome and the ribosome.^6,7^ The molecular machines function by moving their constituting parts, i.e the individual proteins, relative to each other - similar to the way the parts of a mechanical machine move. To enable these fast structural rearrangements, the protein-protein interactions have to be highly specific but also reversible.^8,9^ These contradicting requirements are achieved by a careful balance of electrostatic forces, hydrophobic interactions and hydrogen bonds.^10–12^

Protein domains that serve as a molecular coupling device between proteins have evolved. For example, WW domains selectively recognise proline-rich motifs.^13–16^ A protein that contains a WW domain can selectively bind to another protein that features a matching proline-rich sequence on its surface. WW domains are employed in a wide variety of molecular machines,^17–19^ and to date five different consensus target sequences have been identified.^15,16^ A detailed picture of the relevant binding interactions between single WW domains and their matching proline-rich motifs^16^ emerged from several high-resolution structures of single WW domains bound to peptide ligands. ^20–23^

WW domains can be repeated up to four times in the protein structure, which increases the specificity of the protein-protein interaction.^24^ Tandem repeats of two WW domains (tWW), connected by a linker region of varying length, occur particularly frequently. ^25,26^ tWWs can bind with both WW domains to the same protein and thereby recognize a longer peptide sequence with two proline-rich consensus motifs, or each of the WW domains can bind to a different protein thereby stabilizing a trimeric protein complex.^16,25,26^ In addition to the interactions with the proline-rich sequence, tWW complexes are stabilized by highly specific interactions between the two WW domains. Finally, synergistic effects from the tWW as well as from the ligand can further modulate the binding equilibrium. ^26^ Thus, the structural elucidation of tWW protein complexes is much more demanding than that of single WW domain complexes.

Recently, the crystal structures of four different tWWs in complex with proline-rich peptide ligands have been published.^27–29^ The complex structures of three of these tWWs (KIBRA, MAGI2 and SAV1) show a similar relative orientation of the two WW domains. The two WW domains are in close contact and the complex is stabilized by highly specific interdomain interactions. In the complex of the fourth tWW domain (YAP1), the two WW domains barely touch and assume a spatial arrangement that differs distinctly from the spatial arrangement of the WW domains in KIBRA, MAGI2 and SAV1.

Furthermore, the proline-rich peptide sequences were found to bind in two different orientations to the tWW. In the parallel orientation, the C-terminus of the peptide binds to the C-terminal WW domain and the N-terminus binds to the N-terminal WW domain (YAP1). In the antiparallel orientation, the orientation of the peptide ligand is reversed, such that the N-terminus of the ligand binds to the C-terminal WW domain (KIBRA, MAGI2 and SAV1). However, it is not known what determines the orientation of the ligand.^27–29^

Since the binding equilibrium of tWW domains is influenced by all of these different factors, known complex structures cannot serve as templates for unknown structures of tWW complexes. Rather the complex structure has to be elucidated for each tWW-complex separately. Here we propose a computational workflow to predict tWW complex structures. We use this workflow to predict complex structures of the human formin-binding protein 21 (h-FBP21, Fig. 1.a) to a proline-rich sequence of the core-splicing protein SmB/B’ (Fig. 1.b).

**Figure 1:**
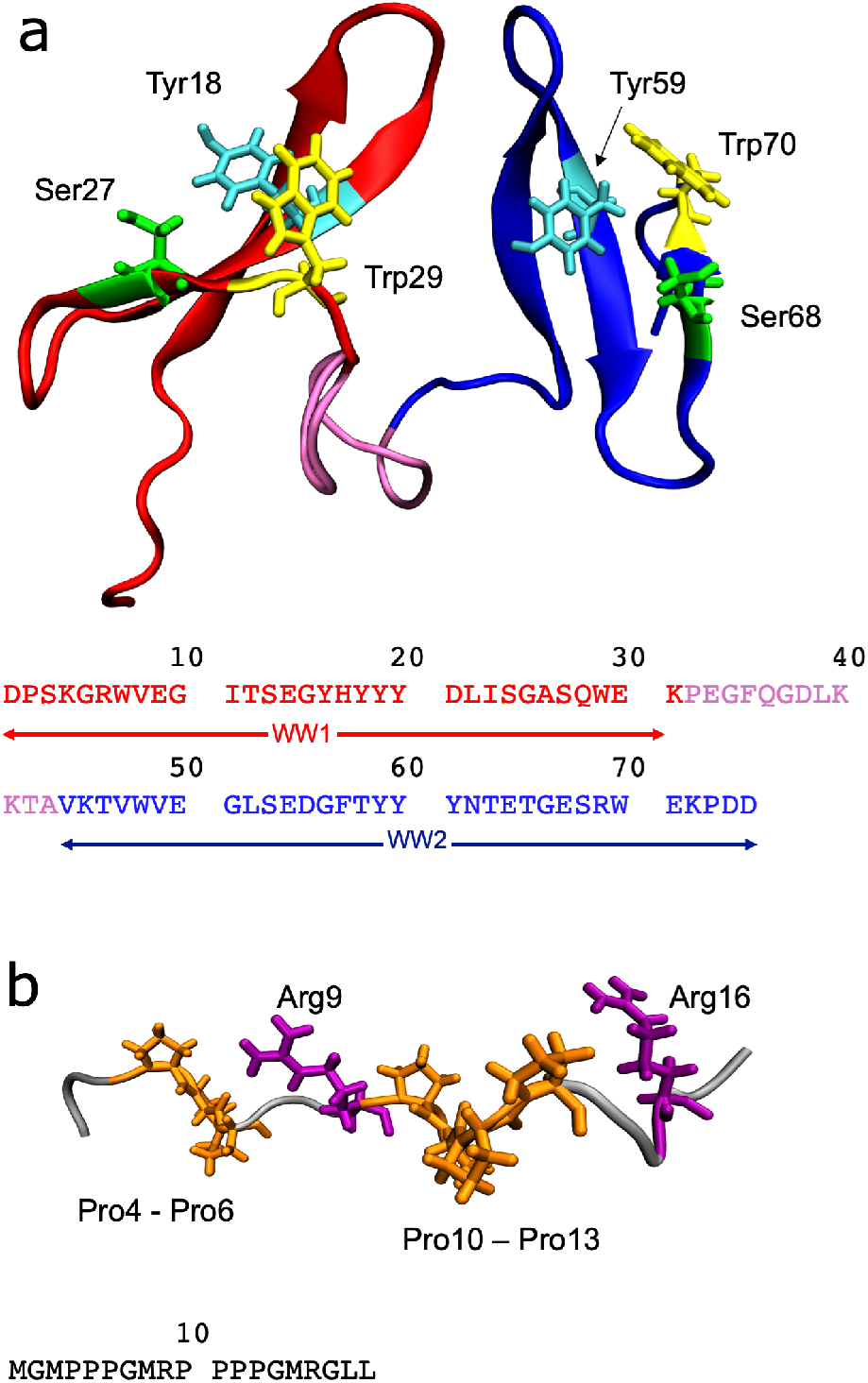
**a**-**b**: Structures and amino acid sequences of the h-FBP21 tWW (**a**) and the SmB2 ligand (**b**). The individual WW domains as well as the interdomain region are coloured as follows. Red: N-terminal WW domain, pink: linker region, blue: C-terminal WW domain. The residues that are critical for binding to the proline-rich motif are shown in the stick representation.

h-FBP21 is part of the spliceosome and is a particularly challenging tWW. The two WW domains are connected by a flexible linker and can move freely with respect to each other in the unbound state. From previous studies we know that h-FBP21 tWW binds peptides with two proline-rich motifs with much higher affinity than peptides with a single proline-rich motif, exceeding the summed affinities of the single domains.^30,31^ When adding a spin-label to the N-terminus of the proline-rich ligand, the spin-label interacted with both WW domains, ^30^ which indicates that the ligand can bind both in the parallel and in the antiparallel orientation to the tWW domain of h-FBP21.

We combine molecular-dynamics simulations and NMR experiments to propose two new structures for the h-FBP21 tWW / SmB/B’ complex. We assume that the binding occurs via conformational selection of a binding-competent apo-conformation, and develop a systematic workflow to identify potentially binding-competent structures of the apo-tWW. From these binding-competent conformations, we generate putative complex structures and assess them with respect to stability and agreement with experimental data. We identify two possible complex structures and characterize them in detail. Our workflow may serve as a blueprint for the computational analysis of protein-protein interactions.

## 2 Results and discussion

### 2.1 Structure of h-FBP21 tWW and SmB2 peptide

#### 2.1.1 General features

The tandem WW domain of h-FBP21 (h-FBP21 tWW) consists of two WW domains, WW1 and WW2, connected by a short, but flexible interdomain region (linker, residues 32-43). The WW domains share 53% sequence identity and show the known fold for WW domains: a triple-stranded, antiparallel *β*-sheet^32^ (Fig 1.a). h-FBP21 tWW recognizes proline-rich sequences, which contain either the motif PPLP (group II motif) or the motif PPR (group III motif).^32^

WW domains recognise proline-rich motifs by forming a highly conserved hydrophobic groove (XP groove) consisting of a Trp residue at the end of the third *β*-strand and a Tyr located in the middle of the second *β*-strand. ^16,20,22^ The *β*-sheet fold positions these two residues opposite of each other, such that their aromatic side chains are perpendicular to each other (Fig 1.a). Two consecutive prolines, P1 and P2 of the proline-rich ligand tightly pack into this hydrophobic groove, where P1 forms a hydrophobic contact with Trp and P2 forms a hydrophobic contact with Tyr. ^16,27,29^ We will refer to these two prolines as the PP motif. In h-FBP21 tWW, the residue pairs are Tyr18 and Trp29 in WW1 and Tyr59 and Trp70 in WW2. Additionally, Tyr20 (WW1) and Tyr61 (WW2) contribute to the hydrophobic interactions. ^30^

The complex is further stabilized by two conserved hydrogen bonds between the WW domain side chains and the ligand backbone. The Trp side chain from the XP groove forms a hydrogen bond to a backbone carbonyl oxygen that precedes the PP motif, and a nearby Ser or Thr forms a hydrogen bond to a backbone carbonyl oxygen that comes after the PP motif in the ligand peptide chain. ^16^ In h-FBP21 tWW, this second hydrogen bond donor is Ser27 in WW1 and Ser68 in WW2 ^20,32^ (Fig 1.a).

WW domains can recognise proline-rich motifs in two opposite orientations, one in which the N-terminal end of the proline-rich motif binds to Trp from the XP groove and the C-terminal end binds to Tyr, and one in which the orientation of the proline-rich motif is reversed.^14^ This gives rise to the parallel and antiparallel orientation of bivalent ligands in tWW domains mentioned in the introduction. However, both ligand orientations are stabilized by the same interactions with the WW domain: hydrophobic contact to Trp/Tyr, hydrogen bonds of backbone carbonyl oxygens with Trp and the nearby Ser.^16^

The ligand in our study (Fig. 1.b) is derived from a proline-rich sequence of the core-splicing protein SmB/B’ (amino acids 213 to 231), and will be denoted SmB2. The “2” indicates that it is a bivalent ligand with two proline-rich motifs (Pro4, Pro5, Pro6 and Pro10, Pro11, Pro12, Pro13). As each proline-rich motif has three consecutive prolines, each motif can in principle bind in two different register shifts to the XP groove, e.g. for the first motif with Pro4 and Pro5 or with Pro5 and Pro6. The SmB2 ligand contains two Arg residues (Arg9 and Arg16), which are separated by two residues from the proline-rich motifs. In WW domains that recognize the PPR motif, the binding affinity and the binding specificity is increased by the Arg residue directly adjacent to the PP motif.^33,34^ Yet, the way the Arg residues in the SmB2 ligand interact with h-FBP21 tWW is not known.

We are now ready to formulate test criteria for putative complex structures. For both WW domains we will test, whether

1. two consecutive prolines are packed into the XP groove (WW1: Tyr18, Trp29, WW2: Tyr59, Trp70)
2. a backbone carbonyl oxygen from the ligand forms a hydrogen bond to the side chain of Trp in the WW domain (WW1: Trp29, WW2: Trp70)
3. a backbone carbonyl oxygen from the ligand forms a hydrogen bond to the side chain of Ser in WW domain (WW1: Ser27, WW2: Ser68)

Additionally, we will analyze the role of Arg9 and Arg16.

#### 2.1.2 Secondary structure of h-FBP21 tWW in the absence of a ligand

We conducted unbiased MD simulations with explicit water (70 independent trajectories, in total simulation length: 35.05 *μ*s) to sample the conformational equilibrium of apo-h-FBP21 tWW. Throughout the simulations, the secondary structures of the two WW domains remain intact. In particular, the *β*-sheets are stable (Fig. 2.a). The loop regions between the *β*-strands occasionally fluctuate between the closely related secondary structure types “bend” and “hydrogen-bonded turn”. By contrast, the linker region between Pro32 and Ala43 is extremely flexible and does not form a stable secondary structure (Fig. 2.a). As a consequence, the two WW domains are only occasionally in direct contact and assume an enormous variety of relative conformations (Fig. 2.b). There is no stable interface between the two WW domains. These findings are in line with NMR/NOE experiments,^22,32^ SAXS experiments,^35^ MD simulations of a single FBP-WW domain^36^ and previous MD simulations of the ligand-free h-FBP21 tWW (apo-h-FBP21 tWW).^35^

**Figure 2:**
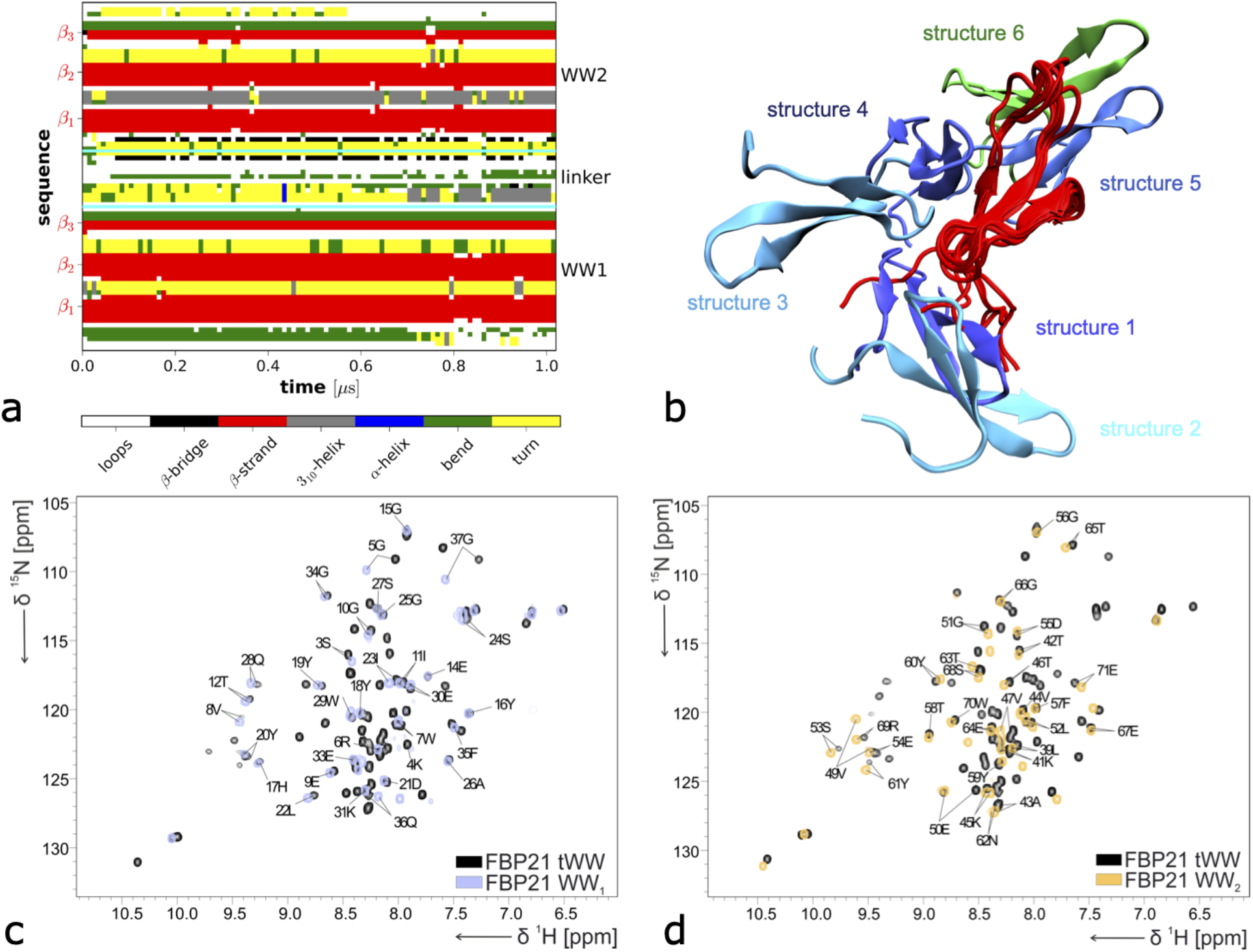
(a) Overlay of six different structures of apo-h-FBP21 tWW resulting from our MD simulations. The structures are aligned on the WW1 domain (red) to highlight the high variability in the relative orientation of the WW2 domain (blue to green). (b) DSSP plot of 1 *μ*s MD simulation of the apo-h-FBP21 tWW. (c,d) Overlay of ^1^H-^15^N-HSQC spectra of the h-FBP21 tWW (black) with isolated WW domain 1 (blue, c) or WW domain 2 (orange, d) in the apo-states.

We further verified the stability of the WW fold in absence of the ligand and in the absence of a stable WW-WW interface by NMR studies. Fig. 2 compares the ^1^H-^15^N-HSQC spectrum of WW1 in the h-FBP21 tWW to the ^1^H-^15^N-HSQC of the single WW domain 1 (c), and likewise the ^1^H-^15^N-HSQC spectra of WW2 in h-FBP21 tWW and the single WW domain 2 (d). As indicated by the large signal dispersion in the spectra, both the tandem repeat as well as the single WW domains of h-FBP21 are folded in the absence of the proline-rich ligand. As apparent from the overlay of the ^1^H-^15^N-HSQC spectra of the h-FBP21 tWW with the single WW domains, we observe only small chemical shifts between them, which clearly shows that the folding of the WW domains is maintained in the tandem repeat and independent of each other.

In accordance with these results, it has been demonstrated that the apo-h-FBP21 tWW does not fold upon binding to the proline-rich peptide^30,32^ - a mechanism that has been observed for other tWWs. ^37^ Instead, we assume the following:

- The relative movement of the two WW domains can essentially be described as the movement of two rigid bodies.
- Because of the flexible linker, the apo-h-FBP21 tWW samples a wide variety of WW-WW interfaces, each with a different relative orientation of the two WW domains.
- At least one of these conformations is a binding competent conformation.
- Binding dominantly occurs via “conformational selection”, i.e. the peptide ligand “selects” this binding competent conformation from the conformational ensemble of the apo-h-FBP21 tWW.

Our strategy therefore is to scan the simulation data set of the apo-h-FBP21 tWW for structures that are potentially competent to bind to SmB2. The next section discusses the workflow in detail.

### 2.2 Identification of complex structures

#### 2.2.1 Conformations with large distances between the XP grooves

The maximal distance between the two proline-rich motifs in the SmB2 ligand is *d*_max_ = 2.65 nm (measured between backbone carbonyl carbons of Pro4 and Pro13). Conformation in which the XP grooves exceed this distance, cannot accommodate the bivalent SmB2 ligand. These extended conformations should be excluded from our sample of apo-conformations before going on to further analyse the sample. To measure the distance between the XP grooves, we chose the outermost residues in the *β*_2_-strands of the two WW domains as reference points: C*_α_*(His17) and C*_α_*(Tyr20) in WW1, and C*_α_*(Thr58) and C*_α_*(Tyr61) in WW2. This defines four interdomain distances: |**r**_17,58_|, |**r**_17,61_|, |**r**_20,58_|, and |**r**_20,61_| (Fig. 3.a). Conformations in which all four distances exceeded *d*_max_ = 2.65 nm were classified as extended conformations and were removed from the sample.

**Figure 3:**
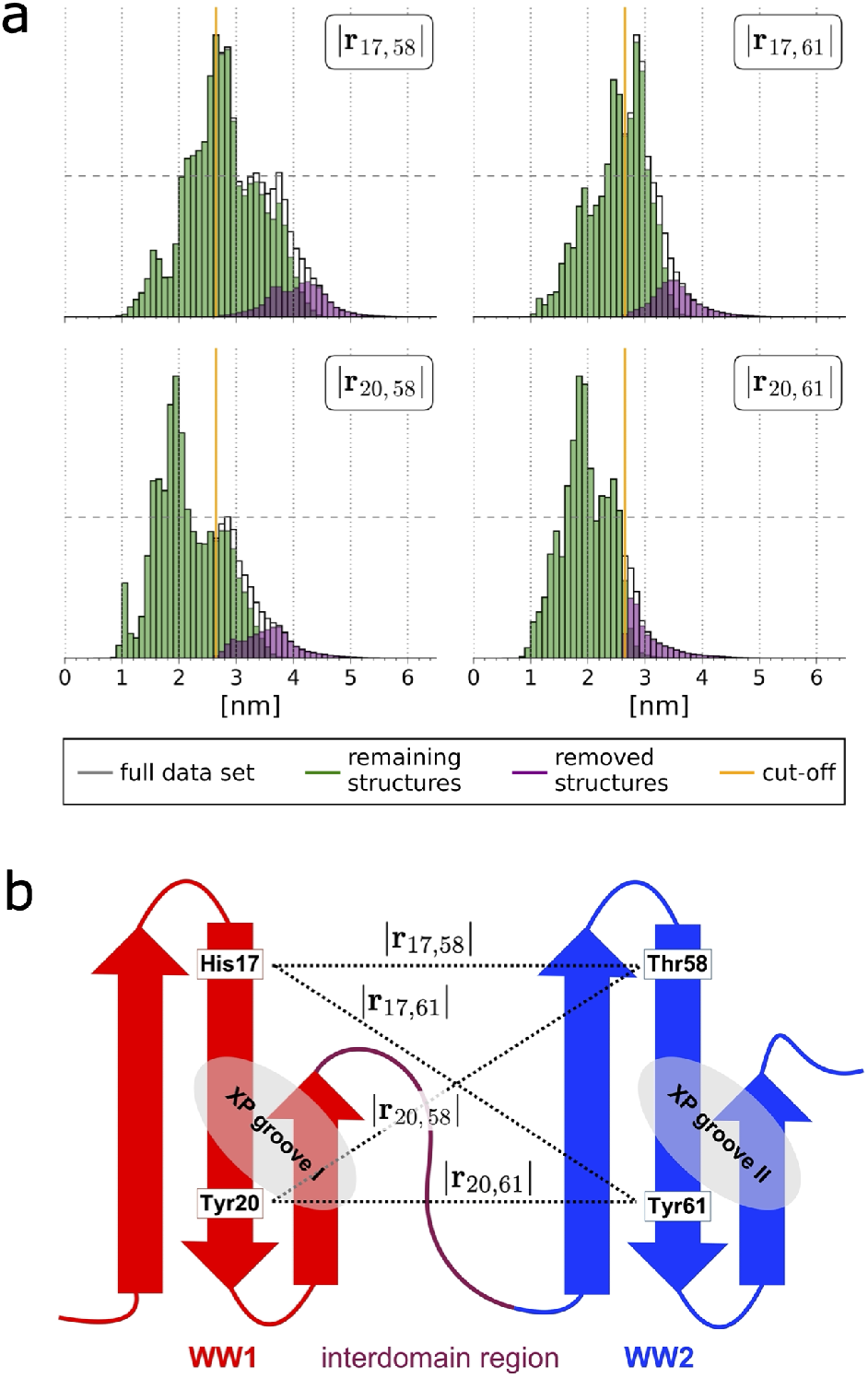
(a) Histograms for the distances |**r**_17,58_|, |**r**_17,61_|, |**r**_20,58_| and |**r**_20,61_|. (b) Location of the reference points in h-FBP21 tWW.

In total, 10.1% of the original sample was removed. Fig. 3.a shows the distance histograms for the original sample (white), the sample of extended conformations (purple), and the sample of compact conformations (green), i.e. after removing the extended conformations. The threshold distance is marked by an orange line. Note that the now reduced sample still contains conformations, in which up to three distances exceed this threshold, and thus the green histograms extend above the threshold line. By contrast, conformations in which all four distances exceeded the threshold were removed, and thus the purple histograms do not extend below the threshold line. The data set of compact conformations contains 31.51 · 10^6^ structures.

#### 2.2.2 Reaction coordinates

To characterise the conformational ensemble in the remaining data set, we projected the data onto a set of reaction coordinates. The two WW domains remain folded and are very rigid, and thus the conformational dynamics of the h-FBP21 tWW can be interpreted as the movement of two rigid bodies. Because of this intuitive interpretation, we devised problem-adapted reaction coordinates rather than relying on data-driven dimensionality reduction techniques. ^38–41^

We used the same four reference points as shown in Fig. 3.b to define the reaction coordinates. Each WW domain is described as a rigid body whose orientation in space is defined by the orientation of its *β*_2_-strand, i.e. by the vector **r**_17,20_ for WW1 and by the vector **r**_58,61_ for WW2. To remove overall translational and rotational motion, we aligned the *β* sheet of WW1 with the *xz*-plane of the coordinate system (Fig. 4.a). For all conformations, this consisted of a translation shift to move C*_α_* (His17) to the origin of the coordinate system, followed by a rotation that aligned **r**_17,20_ with the *z*-axis, followed by a rotation around the *z*-axis that moved C*_α_*(Gly10) into *xz*-plane. The position of WW2 relative to WW1 is now given by the position of Tyr61, i.e. by the vector **r**61 measured in spherical coordinates (*r, θ, ϕ*). The orientation of WW2 relative to WW1 is given by two angles: the torsion angle *α* defined by the four reference points, and the tilt angle *γ* defined by C*_α_*(Thr58), C*_α_*(Tyr61) and C*_α_*(His17) (Fig. 4.a). We did not include the rotation of the *β*-sheet of WW2 around **r**_58,61_ in the set of reaction coordinates. In total, the conformation of the h-FBP21 tWW domain is described by the following five reaction coordinates (*r, θ, ϕ*), *α, γ*.

**Figure 4:**
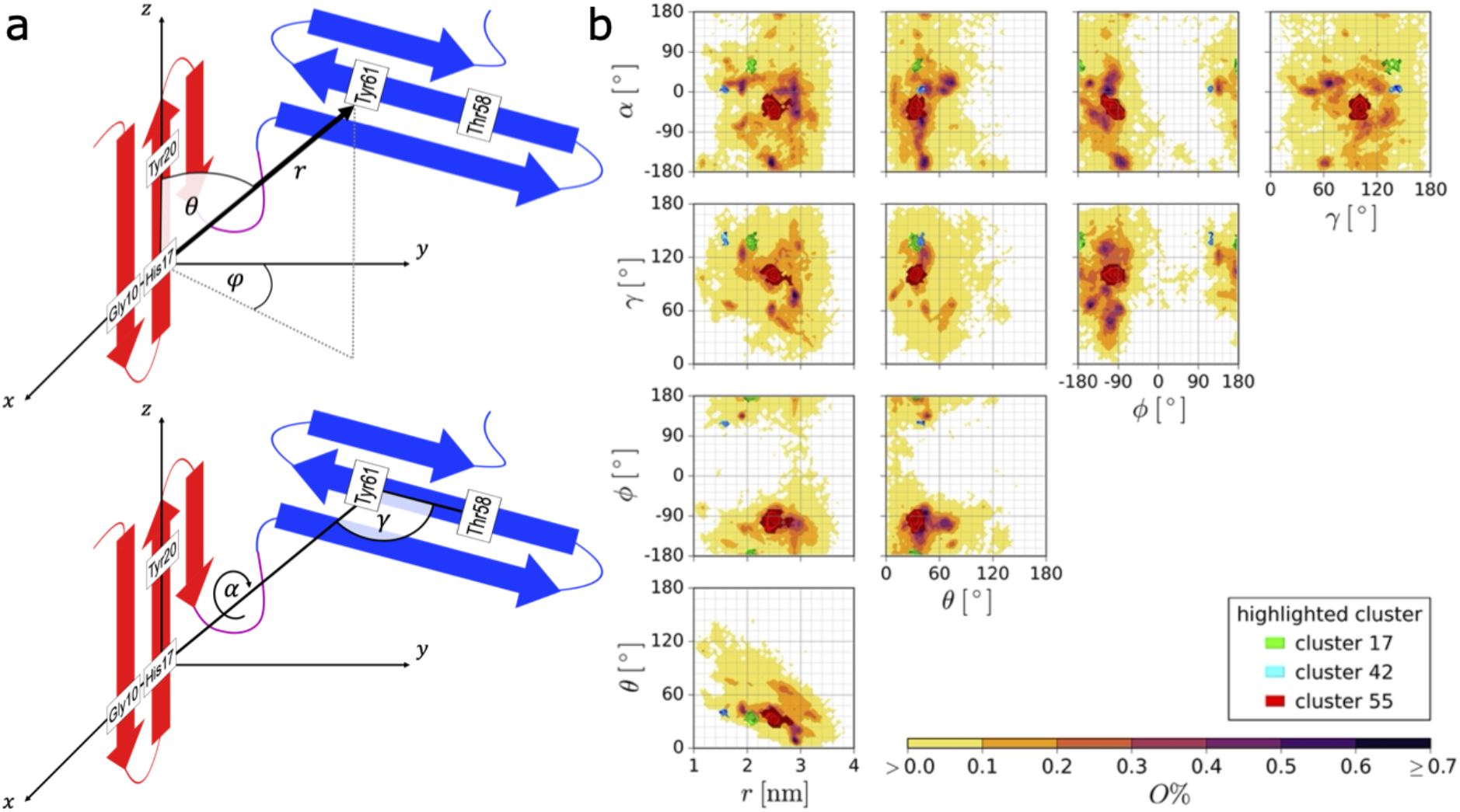
(a) Reaction coordinates (*r, θ, ϕ*), *α, γ*. (b) distribution of the apo-h-FBP21 tWW structures projected onto pairs of reaction coordinates. Each dimension was discretized in 60 equally sized bins. Three clusters (red, green, blue) are highlighted.

Fig. 4.b shows the probability density in this five-dimensional space projected onto pairs of reaction coordinates. The broad probability density with multiple maxima in each of the graphs indicates that the two WW domains move in a shallow and rugged energy landscape. This is in line with the large variety of conformations of the h-FBP21 tWW that we saw in the MD simulation.

#### 2.2.3 Cluster analysis

Next, we extracted highly-populated conformations from the data set of compact structures using the common-nearest-neighbor clustering (CommonNN) algorithm^42–44^ The CommonNN algorithm is a density-based cluster algorithm, which identifies clusters by (approximately) cutting along isodensity lines in the high-dimensional probability density. Subclusters can be identified in a hierarchical fashion by repeatedly applying the clustering algorithm and moving to a higher isodensity line in each hierarchical iteration. Data points in low-probability-density regions of the conformational space, i.e. where the probability density is lower than the lowest isodensity line probed in the hierarchical clustering, are classified as noise points. The CommonNN has been shown to successfully partition challenging test data sets^43^ as well as high-dimensional molecular data sets.^42,44,45^

Prior to clustering, the data set of compact conformations was downsampled to 31, 576 structures. An initial clustering with the CommonNN algorithm and a low density threshold yielded 24 clusters, which were hierarchically refined into 45 clusters comprising 41% of the downsampled data set. The remaining 59% of the data set was classified as noise. The clusters represent the peaks in the probability density, whereas the noise points represent structures in the intermediate regions between the peaks. Fig. 4.b shows the projection of three example clusters in the space of the reaction coordinates.

By construction, the clusters do not overlap in the space of the reaction coordinates. Moreover, the clusters are found in all (populated) regions of the reaction coordinate space (Fig. S1). To judge how well-separated the clusters are in the space of all coordinates, we need to determine whether the structural variation within a cluster is smaller than the structural variation between the clusters. We determined the structural variation RMSD*_ii_* within a cluster *i* as the average RMSD between all pairs of cluster members. We determined the structural variation RMSD*_ij_* between two clusters *i* and *j* as the average RMSD between members of cluster *i* and members of cluster *j*. The procedure essentially amounts to coarse-graining the RMSD matrix of the downsampled data set according to cluster membership. We find that the clusters are rather compact with RMSD*_ii_* varying between 0.15 nm and 0.51 nm (average RMSD*_ii_*: 0.29 nm). By constrast, the structural variation between clusters RMSD*_ij_* varied between 0.38 nm and 1.5 nm (average RMSD*_ij_*: 1.0 nm). In all cases, the structural variation within a given cluster RMSD*_ii_* was smaller than the RMSD*_ij_* to the nearest neighboring cluster (Fig. S2).

Additionally, each cluster has a unique interdomain hydrogen bond pattern (Fig. S3). Interdomain hydrogen bonds are formed between the two WW domains and therefore determine their relative spatial arrangements. Thus, by construction, by the location of the clusters in the space of reaction coordinates, by analysis of the intercluster RMSD and by analysis of the hydrogen bond pattern, the 45 clusters represent distinct conformations.

#### 2.2.4 Protein-peptide docking with HADDOCK2.2 webserver

The 45 clusters represent potentially binding-competent conformations. To judge whether a given conformation of the apo-h-FBP21 tWW domain could indeed form a complex with the SmB2 ligand, we docked the ligand to representative structures of each cluster using the HADDOCK2.2 webserver.^46–48^ The structure of the SmB2 ligand was taken from Ref. 30. For all 45 representative apo-h-FBP21 tWW structures, the docking returned proposals for the h-FBP21 tWW / SmB2 complex. We saved the two proposals with the lowest HADDOCK scores for each tWW structure (i.e. 90 proposals). Each HADDOCK proposal consists of a bundle of four h-FBP21 tWW / SmB2 complex structures. Since the average standard deviation of the HADDOCK score within the proposals (〈σ〉 = 4.49) was sizeable compared to the standard deviation across all h-FBP21 tWW / SmB2 complex structures (σ = 9.43), we did not use the HADDOCK score to further rank the proposals. (See Fig. S4 for an overview of all obtained HADDOCK scores). Instead we used the criteria formulated in section 2.1 to compare the proposals (Table 1).

**Table 1:**
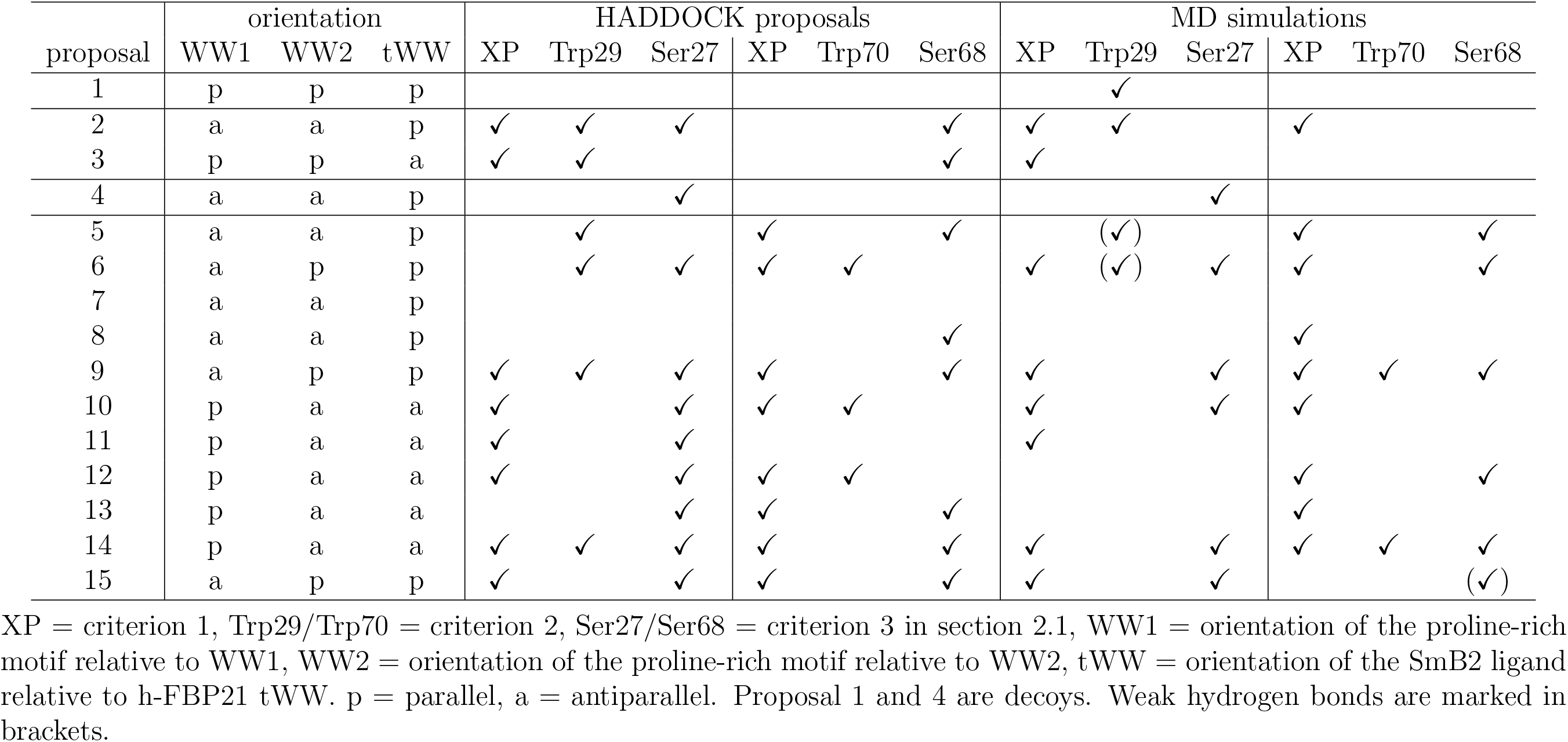
Structural characteristics of the proposals.

13 proposals fulfilled criterion 1 (see section 2.1.1), i.e. the proline-rich motifs of the SmB2 ligand were bound in the vicinity of the XP grooves of the h-FBP21 tWW domain, and were considered for further analysis. In the other 77 proposals, the proline-rich motifs that were not in contact with the XP grooves, were attached elsewhere on the surface of the h-FBP21 tWW. From these, we selected two proposals as decoys for further analysis (proposal 1 and 4). See Fig. S5 for a graphical representation of all 15 selected structures.

The selected proposal structures originated from 11 different CommonNN clusters. (In four clusters, the two HADDOCK proposals with the lowest score both fulfilled criterion 1.) The clusters are located throughout the space of reaction coordinates (Fig. S1), which indicates that many different relative orientations of the two WW domains can potentially accommodate the SmB2 ligand. Table 1 summarizes how the selected 15 proposals fare with respect to the criteria from section 2.1.1.

We remind the reader that the SmB2 ligand can potentially bind in two different orientations to the h-FBP21 tWW: parallel, in which the N-terminus of the ligand binds to the N-terminal WW domain of the h-FBP21 tWW domain and its C-terminus binds to the C-terminal WW domain, and the antiparallel binding mode, in which the orientation of the SmB2 ligand is reversed. The situation is further complicated by the flexible linker. The WW domains can rotate locally, such that their peptide sequence can be oriented either parallel or antiparallel with respect to the peptide sequence of the proline-rich motif in the SmB2 ligand. This gives rise to 8 theoretically possible orientations, which are illustrated in Fig. 5.a. The ligand orientations in the proposal structures are documented in Table 1. Disregarding the decoys, the SmB2 ligand is bound in parallel orientation in 7 out of 13 proposals and in antiparallel orientation in 6 out of 13 proposals. Thus, consistent with experiment, ^30^ the docking calculations do not yield a clearly preferred orientation of the SmB2 ligand. Instead, a dynamic equilibrium between both orientations seems likely.

**Figure 5:**
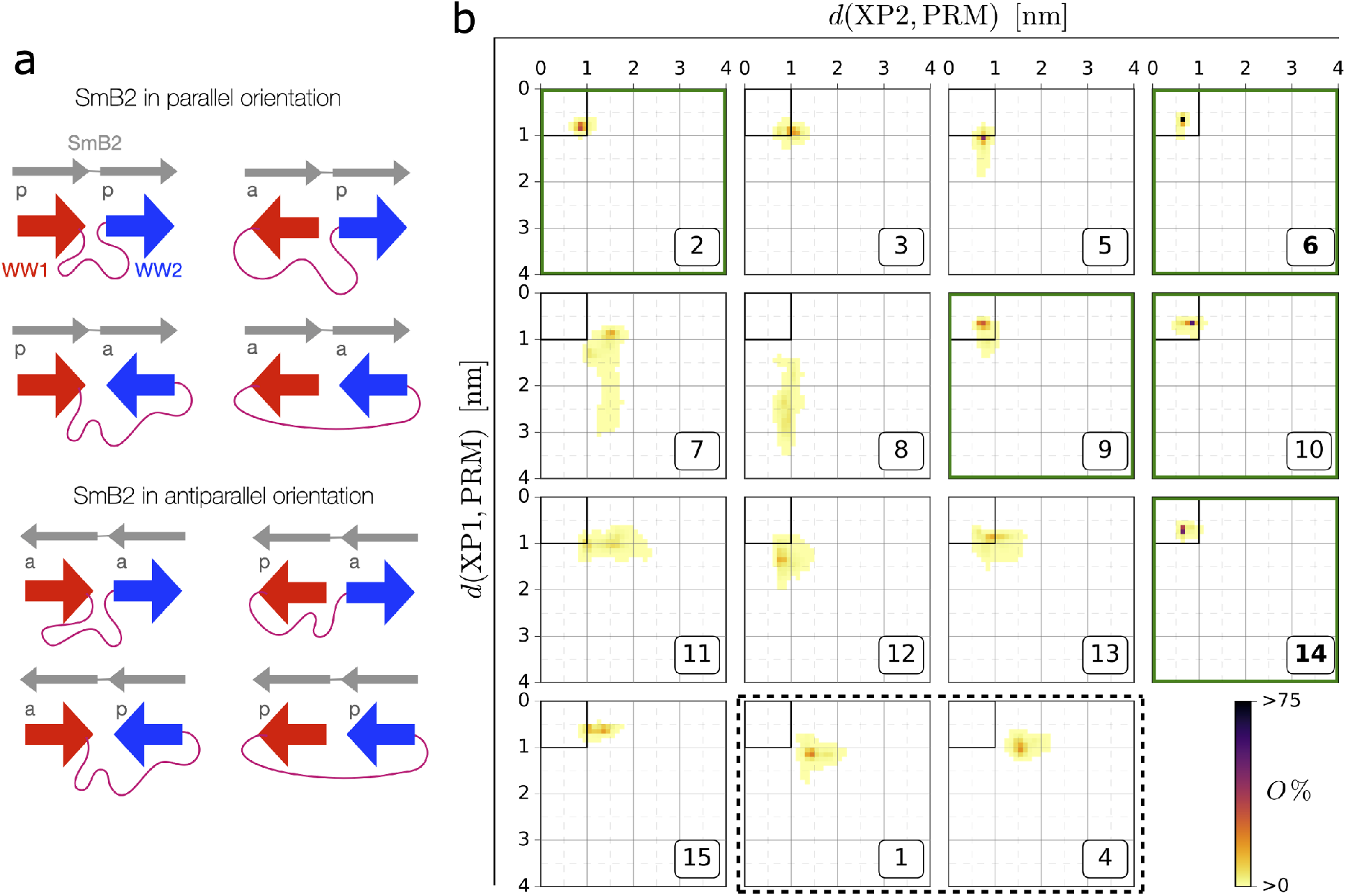
(a) Theoretically possible orientations of SmB2 ligand and tWW. p = parallel, a = antiparallel. (b) Joint probability distributions for the distances between the XP grooves of WW domain 1 (XP1) or 2 (XP2) of the h-FBP21 tWW and their respective closest proline-rich motif of the SmB2 ligand. The target area in which both distances *d*(XP1, PRM) and *d*(XP2, PRM) are smaller than 1 nm is shown as a black square. Green frame: More than 90% of the probability distribution is located in the target area. Dashed black frame: decoys.

To examine the stability of the 15 proposal structures for the complex, we simulated them in explicit water for 50 ns. We gauged whether a complex relaxed into a conformation in which the experimentally known contacts between h-FBP21 tWW and proline-rich ligands are formed, by monitoring the distance between the XP grooves and their respective closest proline-rich motif. We considered the contact formed, if the corresponding distance was below 1 nm. Thus, in the two-dimensional distance distribution plots in Fig. 5.b, a fully formed complex corresponds to the target area marked as black square in the upper left corner of the plots. In some cases, one of the proline-rich motifs diffused away from the XP groove (e.g. 8 and 7, Fig. 5.b), but in none of the 15 h-FBP21 tWW / SmB2 complex structures, the SmB2 ligand fully dissociated from the h-FBP21 tWW. Structures 2, 6, 9, 10 and 14 hit the target area and thus fulfill criterion 1. The remaining proposal structures (1, 3-5, 7,8, 11-13, 15) are discarded.

The discarded structures additionally violated criterion 2 and/or 3. (See Fig. S6 for an overview of the intermolecular hydrogen bonds). Of these structures, only proposal structure 1 has a strong hydrogen bond (>70% population) to Trp29 and thereby meets condition 1. However, since in this structure the SmB2 ligand is clearly not bound to either of the XP grooves, we can safely discard this structure. Structure 5 shows a weak hydrogen bond (<20% population) to Trp29, but the distance of the SmB2 ligand to the XP groove of WW1 is larger than 1 nm, and is thus not consistent with the experimental data. Four of the discarded proposal structures (4, 5, 12, 15) show weak (<40% population) or medium strong (40%-70% population) hydrogen bonds to either Ser27 or Ser68 (criterion 3), but fail the other criteria. Structures 3, 7, 8, 11 and 13 fail all three criteria (Fig. 5.b and Fig. S6). Structures 4 and 1, which we included as decoys, are correctly identified as structures that are not consistent with experimental data and are discarded.

Of the five structures that hit the target area, we discarded structures 2, 9, and 10. Structure 2 shows a highly populated (> 90%) hydrogen bond to Trp29 of the XP1. However, the contact between XP2 and the SmB2 ligand is not stabilized by a hydrogen bond to Trp70, but instead by an ionic bridge between side chains of Asp75 in the h-FBP21 tWW domain and Arg9 in the SmB2 ligand. Thus, criterion 2 is only fulfilled for XP1. The complex does not have any hydrogen bonds to Ser27 or Ser68 (criterion 3). Structure 9 partially samples the hydrogen bonds in criterion 2 and 3. It has the same alignment as 6, and during the simulation 9 relaxed towards 6. Therefore, we only included structure 6 in the further analysis. Structure 10 does not have any hydrogen bond to Trp29 or Trp70 (criterion 2) and is stabilized only by hydrogen bonds to Ser27 and Tyr18. Additionally, visual inspection showed that the proline-rich motifs are poorly positioned in the XP grooves. Overall, 6 and 14 emerged as possible candidates for the complex structure and were analyzed further in section 2.3.

We remark that in 11 of the 15 proposal structures in Fig. 5.b, Arg9 or Arg16 of the SmB2 ligand form an ionic bridge to residues in the h-FBP21 tWW domain (See Fig. S6 for intermolecular hydrogen bonds). We investigate this finding further in section 2.3.4.

### 2.3 Complex structures 6 and 14

To characterize the two candidate structures, we extended the MD simulations of the complexes 6 and 14 to 0.5 *μ*s each. In this section we discuss the interface between the two WW domains in both complex structures. In the next section, we discuss the interactions between h-FBP21 tWW and the SmB2 ligand. The coordinate files of structures of 6 and 14 have been published on Zenodo (https://doi.org/10.5281/zenodo.5680225).

#### 2.3.1 The WW1-WW2 interface

Both complexes remained stable throughout the simulation, i.e. the two proline rich motifs of the SmB2 ligand remained tightly bound to the XP grooves of h-FBP21 tWW (Fig. S7.a), and the WW domains retained their relative orientation (Fig. S7.b). Thus, h-FBP21 tWW does not relax towards a structure which is capable of binding the ligand in both orientations.

Even though the single WW-domains can bind to a single proline-rich motif in parallel or antiparallel orientation, the tandem-WW domain discriminates between the orientations of the bivalent SmB2 ligand. It adopts distinct binding-competent structures for the parallel and for the antiparallel orientation of SmB2.

Fig. 6.a shows the different orientations of WW1 relative to WW2 in the complex structures 6 and 14. There is a relatively large gap between WW1 and WW2 in complex structure 6, whereas the WW domains are in direct contact in structure 14. This is in line with the observation that structure 6 is more flexible than 14, as indicated by the broader distribution of 6 in Fig. S7.b.

**Figure 6:**
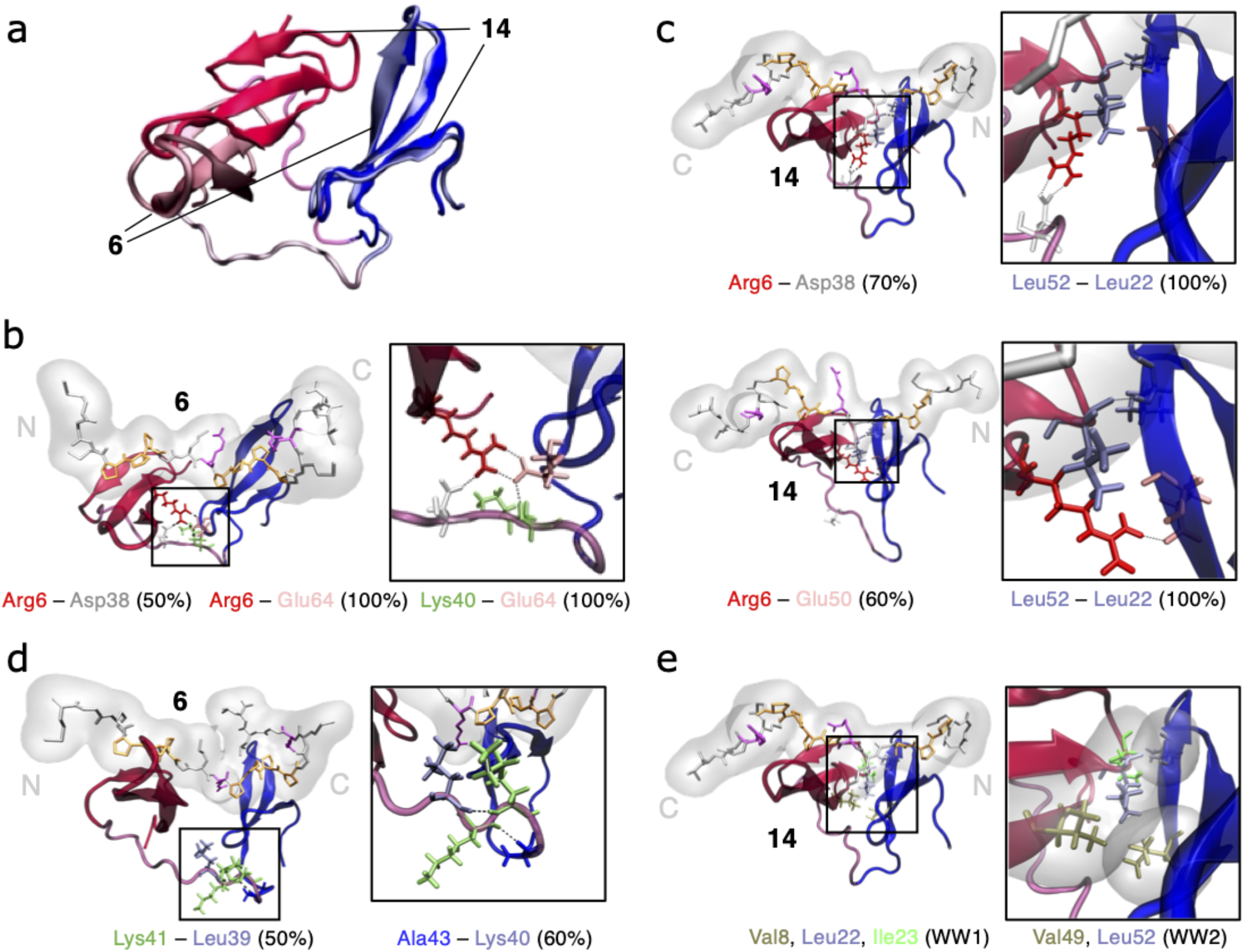
(a) Overlay of the h-FBP21 tWW structures in 6 and 14. The structures are aligned on WW domain 2 (blue). (b,c) Interdomain hydrogen bonds for the complexes 6 (b) and 14 (c) with populations as indicated. Further structure-determining interactions, such as intradomain hydrogen bonds in complex 6 (d) and hydrophobic contacts in complex 14 (e) are shown. The involved residues are coloured according to the residue type. Red: arginine, white: aspartate, light blue: leucine, pink: glutamate, green: lysine, blue: alanine, ochre: valine.

In structure 6, the domain-domain interface is stabilized by an intricate hydrogen bond network that heavily relies on residues in the linker. The positively charged side chain of Arg6 in the N-terminal tail of WW1 forms a strong ionic bridge to the carboxyl group of Glu64 in the loop region between the *β*_1_- and *β*_2_-strands of WW2. Additionally, the carboxyl group of Glu64 forms a strong hydrogen bond to the backbone amide of Lys40, and the side chain of Arg6 forms a somewhat weaker hydrogen bond to the carboxyl group of Asp38 (Fig. 6.b). In our simulations, we find conformations with a slightly twisted linker, where the twist is stabilized by a hydrogen bond between backbone amide of Lys41 and the backbone carbonyl group of Leu39, as well as a hydrogen bond between the backbone amide of Ala43 and the backbone carbonyl group of Lys40 (Fig. 6.d). There are no hydrophobic interactions that stabilize structure 6.

By contrast, structure 14 is predominantly stabilized by a hydrophobic contact between the two WW domains, involving Val8, Leu22 and Ile23 in WW1, and Val49 and Leu62 in WW2 (Fig. 6.e). The hydrophobic contact is further stabilized by a strong hydrogen bond between the backbone amide of Leu52 and the backbone carbonyl group of Leu22 (Fig. 6.c). However, also Arg6 plays a role. It can switch between two conformations: one in which it forms an ionic bridge to the carboxyl group of Asp38, and one in which it forms a hydrogen bond to the backbone amide of Glu50 (Fig. 6.c).

#### 2.3.2 The interactions between h-FBP21 tWW and SmB2 ligand

In complex 6, the SmB2 ligand has a parallel orientation, i.e. its N-terminal proline-rich motif binds to the N-terminal WW1 domain, and its C-terminal proline-rich motif binds to the C-terminal WW2 domain. As expected, both proline-rich motifs are tightly packed into the XP grooves formed by Tyr18 and Trp29 in WW1, and by Tyr59 and Trp70 in WW2. The N-terminal proline-rich motif is additionally stabilized by the literature-known hydrogen bonds from Trp29 and Ser27 to backbone carbonyl oxygens in the ligand (Fig. 7.a) Thus, the interaction between the N-terminal proline-rich motif and WW1 shows all the hallmarks of a typical complex between a WW domain and a proline-rich sequence. Additionally, the Smb2 ligand has two methionine residues (Met1, Met3) just before the N-terminal proline-rich motif. This generates an extended hydrophobic surface, which wraps around Trp29 in the WW1 domain (Fig. 7.a). The C-terminal proline-rich motif is also stabilized by the literature-known hydrogen bonds from Trp59 and Ser68 to backbone carbonyl oxygens in the ligand. Additionally, a hydrogen between the side chain of Arg16 in the SmB2 ligand and Glu54 at the turn of the *β*-sheet in WW2 further anchors the ligand to the h-FBP21 tWW domain (Fig. 7.a). The SmB2 ligand can fold onto itself and form a hydrophobic contact between its Leu18 and Leu52 in the *β*-sheet of WW2 (Fig. S8.a).

**Figure 7:**
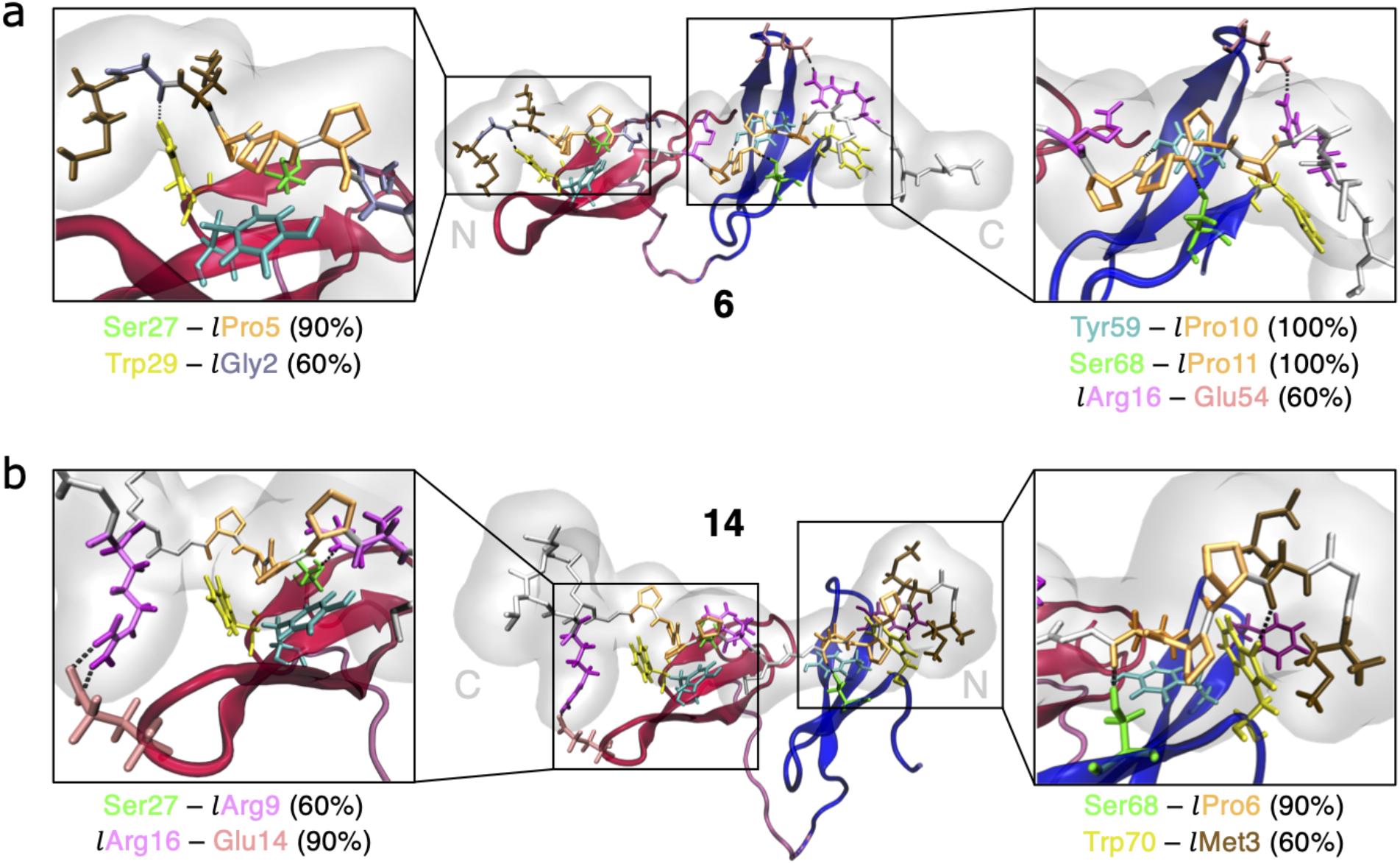
Intermolecular hydrogen bonds for the complexes 6 (a) and 14 (b) with populations indicated. All residues involved are highlighted according to residue type: tryptophan: yellow, serine: green, tyrosine: cyan, arginine: magenta, proline: orange, glycine: light blue, glutamate: pink, methionine: brown. Residues belonging to the SmB2 ligand are labelled with ‘l’. Residues not involved but important for the interaction between SmB2 ligand and h-FBP21 tWW domain are highlighted according to residue type as well: phenylalanine: purple, methionine: brown.

In complex 14, the SmB2 ligand has an antiparallel orientation, i.e. its C-terminal proline-rich motif binds to WW1, and its N-terminal proline-rich motif binds to WW2. Both proline-rich motifs are tightly packed into the two XP grooves (Fig. 7.b, S8.b). The N-terminal proline-rich motif is stabilized in the XP groove of WW2 by the literature-known hydrogen bonds from Trp70 and Ser68 in WW2 to the backbone carbonyl oxygens of SmB2. In this particular binding interface, Trp70 is entirely shielded from the solvent by a hydrophobic pocket formed by residues preceding the proline-rich sequence, Met1 and Met3 in SmB2, as well as Phe57 in WW2. By contrast, the C-terminal proline-rich motif is not stabilized by the typical hydrogen bonds involving Tyr18 or Trp29 in the XP groove of WW1. We detect a hydrogen bond between Ser27 and Pro10 in the SmB2 ligand, but its population is low, and Ser27 seems to fluctuate between this hydrogen bond and a strong hydrogen bond to Arg9 in the SmB2 ligand (Fig. 7.b). The SmB2 ligand is further anchored in the XP groove by an ionic bridge between Arg16 in the SmB2 ligand and Glu14 in the h-FBP21 tWW, located in the loop region between *β*_1_- and *β*_2_-strands of the WW1 *β*-sheet (Fig. 7.b). Additionally, Met8 in the SmB2 ligand forms a hydrophobic contact with Val8 in WW1 and thereby expands the hydrophobic surface at the interface of WW1 and WW2 (Fig. S8.b) We characterized h-FBP21 tWW in complex with the SmB2 ligand by ^1^H-^15^N-NOESY-HSQC spectra (Table S1). All detectable NOEs between backbone amides were either within WW1 or within WW2 confirming that the *β*-sheets of WW1 and WW2 remain stable in the complex - in agreement with our simulations. More importantly, we could not detect any NOEs between the WW domains. This suggests that the two WW domains are not in stable, direct contact in the h-FBP21 tWW / SmB2 complex - supporting complex structure 6. The NOE data do not rule out complex structure 14, but it indicates that this structure is not the most populated complex structure.

To identify the residues affected most by the binding process, we measured ^1^H-^15^N-HSQC spectra for the spectra for h-FBP21 tWW in complex with the SmB2 ligand (holo-h-FBP21) using saturating ligand concentrations (40 fold molar excess) and determined the chemical shift differences to the apo-h-FBP21 tWW (Fig. S8.c). 10 residues show chemical shift differences above the average plus standard deviation and are therefore considered significantly affected by the complex formation: His17, Tyr18, Trp29 and Glu30 in WW domain 1 and Ser53, Glu54, Tyr59, Tyr60, Trp70 and Glu71 in WW domain 2. Except for Ser53 and Glu54, all these residues are direct members of the XP grooves of WW domain 1 (Tyr18, Trp29) and WW domain 2 (Tyr59, Trp70) or neighbouring residues, which are affected by the environmental changes. The chemical shift differences in Ser53 and Glu54 might be caused by the interaction with the Arg16 of the SmB2 ligand in complex 6. We do not find a chemical shift difference that could be attributed to the analogous hydrogen bond between Arg16 and Glu14 in complex 14 (Fig. 7.b). Again, this finding does not rule out complex 14, but indicates that it is not the most populated complex structure.

#### 2.3.3 Does Arg6 in WW1 stabilize the complex?

Our two proposals for the complex structure differ in interactions formed by Arg6 in the N-terminal tail of WW domain 1. Additionally, Arg6 forms a multitude of hydrogen bonds and ionic bridges to residues in the interdomain region and in the WW domain 2 in the 42 potentially binding competent clusters of apo-h-FBP21 tWW (Fig. 8.a). Interaction partners for ionic bridges are the negatively charged side chains of aspartate (Aps38, Asp74, Aps75) and glutamate (Glu50, Glu64), which are mostly located in WW2. But we also find hydrogen bonds to neutral side chains (Gln36) and backbone carbonyl atoms (Leu39, Pro73). In most cases, the hydrogen bonds or ionic bridges are very strong with populations between 90% and 100%. Even though Arg6 is typically not part of the immediate WW1-WW2 interface, its interaction with WW2 seems to be characteristic for each of the clusters (Fig. 8.a).

**Figure 8:**
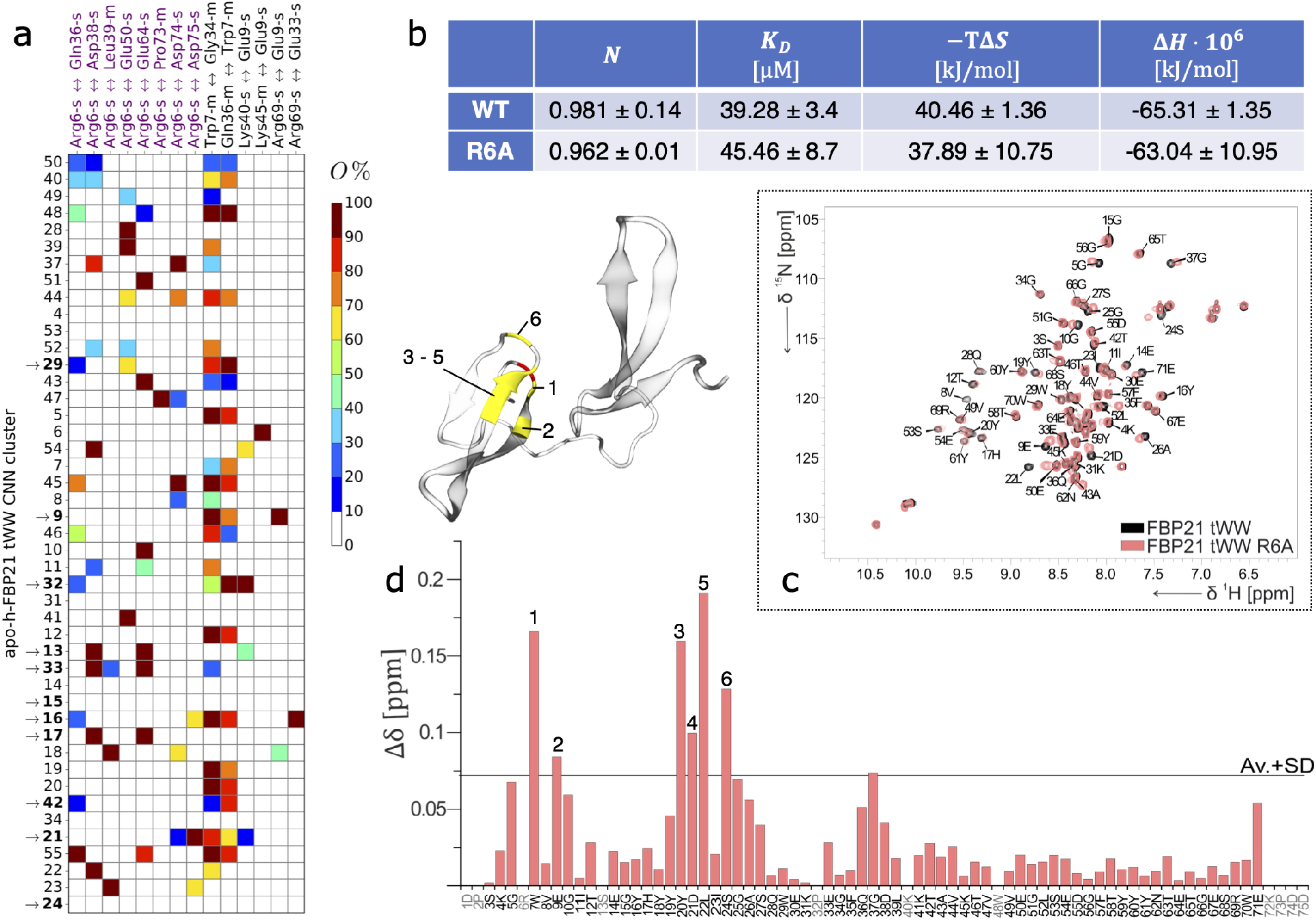
(a) Interdomain hydrogen bonds and ionic bridges denoted as “donor↔acceptor”, that occur at least in one cluster with a relative occurrence *O* > 99%. Clusters sorted by size in decreasing order (top to bottom) and labelled with ‘→’, if the derived complex structure from HADDOCK was simulated. Hydrogen bonds involving Arg6 are highlighted in purple. (b) Results of the ITC experiments for the wild-type h-FBP21 tWW (‘WT’) and its mutant tWW-R6A (‘R6A’) with the SmB2 ligand respectively. Number of bound ligands (*N*), dissociation constants (KD), temperature times entropy changes (-TΔS) and changes in enthalpy (ΔH) are given as average values with standard deviations. See Fig. S9. for the ITC assay (c) Overlay of the ^1^H-^15^N-HSQC spectra of apo-h-FBP21 tWW (black) and the apo-h-FBP21 tWW-R6A (pink). (d) ^1^H-^15^N chemical shift differences between the h-FBP21 tWW and its mutant tWW-R6A. Differences larger than the average plus standard deviation (Av.+SD, black line) are considered significant and labelled with numbers: Trp7 (1), Glu9 (2), Tyr20 (3), Asp21 (4), Leu22 (5) and Ser24 (6) These residues and the mutation site are highlighted in yellow and red, respectively, in the example structure.

Is the specificity of the Arg6 interaction a by-product of a given interface between WW1 and WW2, where Arg6 in the flexible N-terminal tail seeks the nearest negative charge and sticks to it, or is the Arg6 interaction with WW2 critical for the stability of the complex with SmB2? To investigate this question, we substituted Arg6 by alanine in vitro and characterized the mutated protein (tWW-R6A) by NMR and Isothermal Titration Calorimetry (ITC) experiments. In the overlay of the ^1^H-^15^N-HSQC spectra of the apo-h-FBP21 tWW and its mutant tWW-R6A (Fig. 8.c), we observe a close match, which indicates that the overall structure of the h-FBP21 tWW-R6A is maintained despite the mutation. Analysing the chemical shift differences between apo-h-FBP21 tWW and its mutant (Fig. 8.d), we detect the most significant differences, i.e. differences larger than the average plus standard deviation, solely for residues in close proximity to the mutation. These residues are part of WW domain 1 in the *β*_1_-strand (Trp7, Glu9), the *β*_2_-strand (Tyr20, Asp21, Leu22) and in the loop region between the *β*_2_- and the *β*_3_-strand (Ser24). Besides the expected, most affected residues, Gly37 located in the interdomain region, also shows a chemical shift slightly exceeding our significance cut-off. As this residue, however, is located in the fully-flexible part of the tWW, we can deduce that Arg6 does not massively influence the conformation of apo-h-FBP21 tWW. Thus, the secondary structure of apo-h-FBP21 tWW is not affected by the substitution.

In order to rule out that the R6A mutation impairs the interaction with the ligand, we measured the binding affinity of the wild type (WT) h-FBP21 tWW and its mutant tWW-R6A for the SmB2 ligand by ITC experiments. There is no significant difference between the binding affinity of the WT h-FBP21 tWW and its mutant tWW-R6A, measured as dissociation constant (Fig. 8.b). Furthermore, we did not detect any significant differences in the entropic and the enthalpic contributions to the free energy of binding. Thus, Arg6 is not critical for the stability of the complex, and its specific interactions are likely a consequence of an already-existing protein-protein interface between the two WW domains.

#### 2.3.4 The influence of the arginine residues of SmB2 on the interaction with h-FBP21 tWW

In both complexes, the proline-rich motif at the C-terminus of the SmB2 ligand is stabilized in the XP groove by an ionic bridge from Arg16 in the SmB2 ligand to a glutamic acid in the vicinity of the XP groove. Additionally, Arg9 in the SmB2 ligand forms a hydrogen bond to Ser27 in the XP groove of WW1 in complex 14. Arginine residues that flank the proline-rich motif are known to increase the specificity of some ligands,^16^ However, in other cases methylation of flanking arginine residues did not affect the binding affinity. ^34^

We therefore tested the role of Arg9 and Arg16 by mutating them to alanine residues in both h-FBP21 tWW / SmB2 complexes in silico. SmB2-WT denotes the wild-type SmB2 ligand, the single mutants are denoted SmB2-R9A and SmB2-R16A, and the double mutant is denoted SmB2-R9A/R16A. For each mutated complex, we performed 0.5 *μ*s of MD simulations.

Complex 14 does not seem to be affected by the mutation. For all four ligands, the complex remained stable during 0.5 *μ*s of simulation (Fig. 9.a). The interactions between the N-terminal proline-rich motif and the XP groove of WW2 remained stable. The interactions between the C-terminal proline-rich motif and the XP groove of WW1 were most affected by the mutation of Arg9 (green bars in Fig. 9.b), i.e. some critical hydrogen bonds increased in strength while others became less populated. For example, in the complex with the wildtype SmB2 ligand, Ser27 in WW1 fluctuates between a hydrogen bond to the backbone of Arg9 and a hydrogen bond to the backbone of Pro10. Mutating Arg9 destabilizes the first hydrogen bond and instead stabilizes the second. It seems that mutating Arg9 caused a shift in the hydrogen bond patterns in the XP groove of WW1, but overall, the proline-rich motif remained tightly bound. Interestingly, the mutation of Arg9 decreased the strength of the ionic bridge between Arg16 and the glutamate in the h-FBP21 tWW domain (Glu54). The mutation of Arg16 had a less pronounced effect on the hydrogen bond pattern. The effect of the double mutant is somewhere in between the two single mutants.

**Figure 9:**
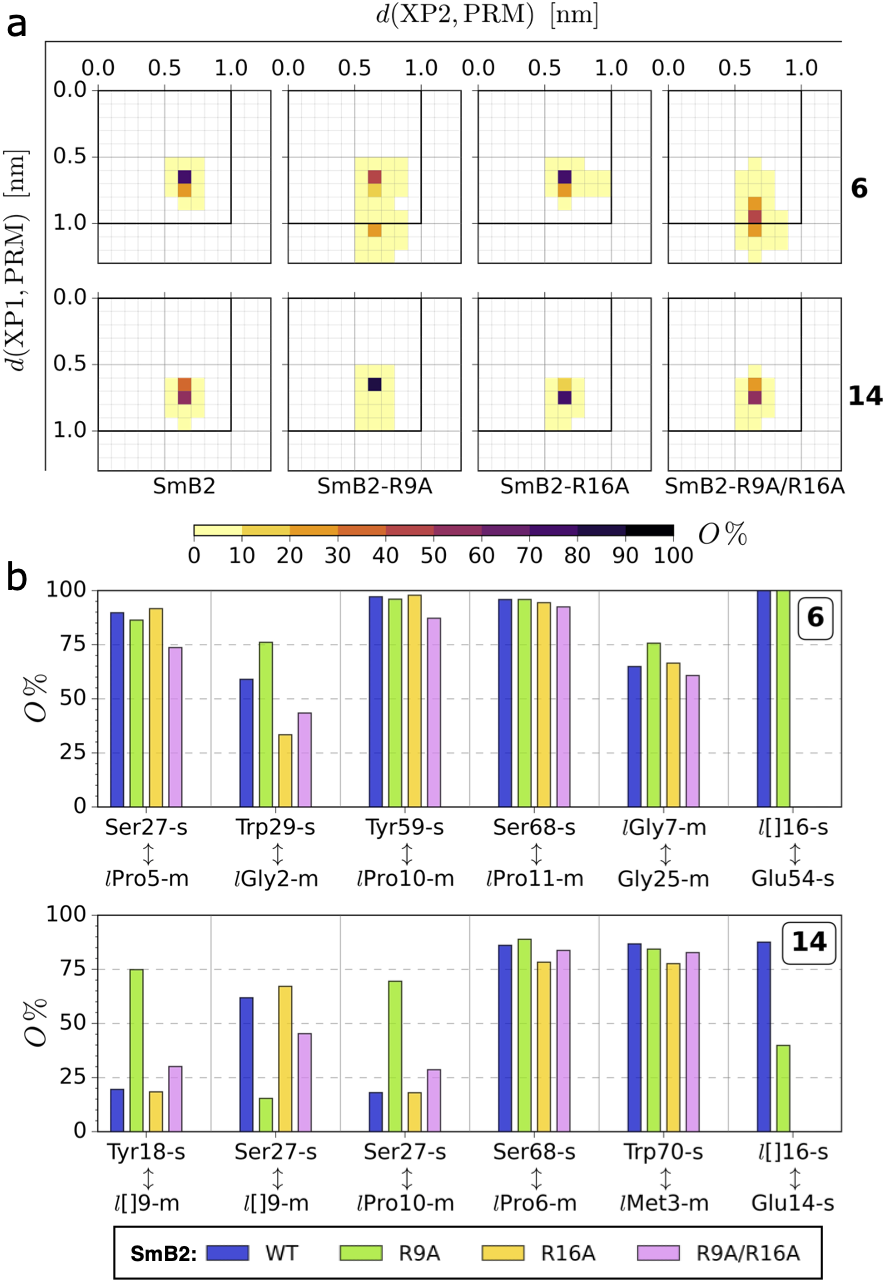
(a) Joint probability distributions for the distances between the XP grooves of the h-FBP21 tWW and the respective closest proline-rich motifs of the (mutated) SmB2 ligand. The bins are equally sized in both distances. (b) Populations on average for inter-molecular hydrogen bonds in the complexes 6 and 14 with a population > 50% in at least one complex. The populations of the hydrogen bonds are presented for each mutation of the SmB2 ligand: blue: wild-type SmB2, green: SmB2-R9A, yellow: SmB2-R16A, purple: SmB2-R9A/R16A.

In complex 6, the C-terminal proline-rich motif is bound to WW2. When we mutated Arg9 (both single and double mutant), we observed that the ligand partially moved out of one XP groove in some of the simulations. However, in each of these cases, the N-terminal proline-rich motif detached from WW1 and not the C-terminal proline-rich motif where the mutated residue resides (Fig. 9.a). We did not observe that the ligand fully detached from the tWW. It is not clear, whether the mutation of Arg9 destabilized complex 6, or whether complex 6 fluctuates more than complex 14, and it was by chance that we observed these fluctuations in the simulations where Arg9 was mutated. Note that mutating Arg9 does not decrease the stability of the critical hydrogen bonds (Fig. 9.b). The strength of the hydrogen bond between Trp29 and the backbone of the SmB2 ligand is even increased. By constrast, mutating Arg16 decreases the strength of this hydrogen bond (Fig. 9.b).

Even though we cannot draw any final conclusion as to thermodynamic stability of the complexes with the mutated SmB2 ligands, the simulations suggest that Arg16 and Arg9 are not decisive for the stability of the two complexes. Complexes 6 and 14 seem to adapt to the mutations without loosing their ability to bind the SmB2 ligands. It is possible that Arg16 and Arg9 aid in the specificity when competing ligands with proline-rich motifs are present, rather than increasing the baseline-stability of the binding.

#### 2.3.5 Induced fit versus conformational selection

Binding of complicated ligands to proteins occurs via a combination of conformational selection- and induced fit-mechanisms. Bivalent ligands, such as SmB2, typically bind by first forming a monovalent contact, followed by a distinct second step in which the second contact is formed and the two binding partners relax into the final complex conformation. Compared to other tWW domains,^26,27,37^ the linker of h-FBP21 tWW is extremely flexible and allows for an enormous variety of relative orientations of the two WW domains in the absence of a ligand. By contrast, the two prolin-rich motifs in the SmB2 ligand are separated by only three residues. It is thus very unlikely that the ligand recruits the second WW domain by an induced fit mechanism. For the ligand to fully bind to both WW domains, the WW domains have to be in approximately the same relative orientation as in the final complex. Thus, the formation of the initial encounter complex in h-FBP21 tWW should be dominated by conformational selection. This has been the underlying assumption for our workflow to identify the complex structures 6 and 14. After the formation of this initial encounter complex, the complexes are stabilized by interactions between h-FBP21 tWW and the SmB2 ligand. Accommodating these interactions likely occurs via induced fit. By comparing the apo-clusters to the final complex structures, we can speculate on the extent to which induced fit plays a role in the binding mechanism.

Overall, the complexes 6 and 14 are similar to their corresponding apo-clusters. However, in both cases we find slight adjustments in the relative orientation of the two WW domains, indicating that induced fit does play a role in the binding process (Fig. 10). Fig. S7.b shows the shifts in the space of reaction coordinates from apo-cluster to complex. In both complexes, a slight shift along *r* occurs, while *θ, γ* and *α* remain unchanged. Complex 14 additionally shows a rotation of about 60° around *ϕ* and a rotation of about 10° in *γ* (Fig. S7.b). This can be related to the binding process. A change in r allows h-FBP21 tWW to adjust to the correct distance between the two proline-rich motifs. In complex 6, where this is the only adjustment, the hydrogen-bond pattern between the two WW domains does not change from apo-cluster to complex (Fig. S3, Fig. S10).

**Figure 10:**
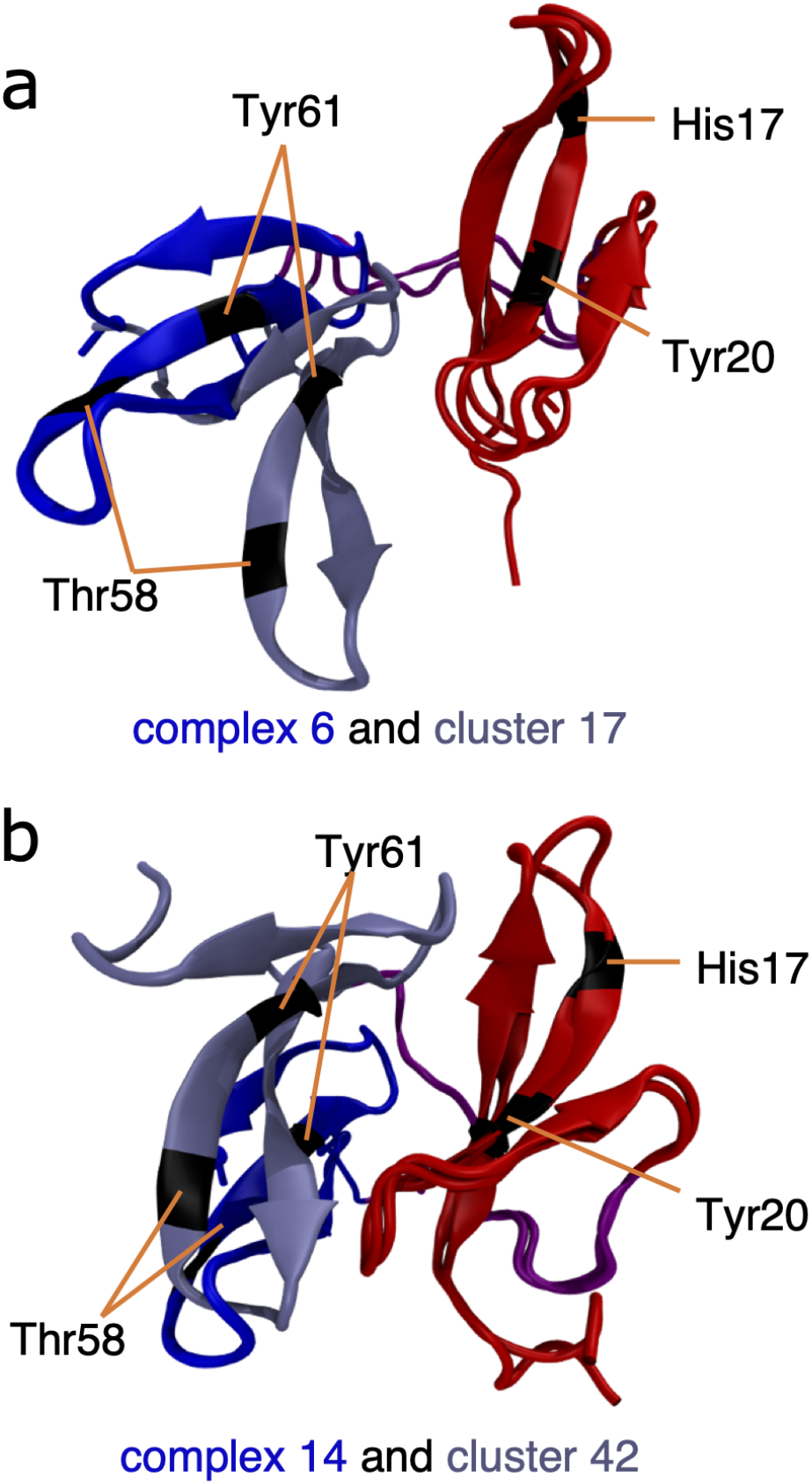
h-FBP21 tWW structures for (a) the CommonNN cluster 17 and the h-FBP21 tWW / SmB2 complex 6 and (b) the CommonNN cluster 42 and h-FBP21 tWW / SmB2 complex 14. The h-FBP21 tWW structures are aligned on the WW domain 1 (red). The WW domain 2 is coloured according to the structure it is originating from: blue (h-FBP21 tWW / SmB2 complex) and pale blue (CommonNN cluster). Residues that were used for the definition of the reaction coordinates are highlighted in black.

For a simultaneous recognition of both proline-rich motifs, the *β*-sheets of the h-FBP21 tWW have to adopt an approximately planar arrangement such that the XP grooves face in the same direction. This is not the case in the apo-cluster of complex 14. In this cluster, the *β*_2_-strands of the WW domains are facing each other (Fig. 10.b), yielding a compact structure. This arrangement is adjusted such that the two XP grooves face in the same direction by a rotation of the WW1 *β*-sheet around its central axis, i.e. by a rotation around *ϕ*, and by the small rotation around *γ*. As a consequence, the interdomain hydrogen bond pattern changes from the apo-cluster to the complex. While interactions between the *β*_2_-strands of WW1 and WW2 are weakened or vanish, the hydrogen bonds between Leu22 and Leu52 as well as the hydrogen bond between Arg6 and Glu50 become more populated (Fig. S3, Fig. S10).

## 3 Methods

### 3.1 Computational methods

All-atom, classical molecular-dynamics simulations were performed with GROMACS simulation packages 5.0.2^49^ and 2016.5^50^ as well as with GROMACS 2019 release version.^51^ The systems were simulated in the *NVT*- and *NpT*-ensembles (1.0 bar, 300 K) with AMBER99SB*-ILDNP^52^ parameters and the TIP3P water model.^53^ Prior to production, starting structures were put into a sufficiently large simulation box, solvated, neutralised and equilibrated for several hundred picoseconds. Before clustering of the apo-h-FBP21 tWW MD data set, structural outliers were removed (see SI 2.2.2 and section 2.2.2) and reaction coordinates were calculated (SI 2.2.3) after ensuring the same orientation of the *β*-sheet of WW domain 1 in all frames. The data set for clustering was defined on chronologically every 1000th frame of the remaining structures expressed in the reaction coordinates. Hierarchical clustering with common-nearest-neighbor clustering (CommonNN)^42–44^ cluster algorithm yielded 45 clusters for the apo-h-FBP21 tWW domain. Docking experiments were performed with HADDOCK2.2 webserver^48^ considering a representative structure for each cluster (see SI 2.2.4) and a structure of the SmB2 ligand,^30^ respectively. The following analyses were performed using built-in functions of the GROMACS software package: Hydrogen bonds were analysed with gmx hbond (see SI 2.2.6 and SI 2.4.1) and the RMSD between a reference structure and another data set was calculated with gmx rms (see SI 2.2.5).^51^ The python library mdtraj 1.9.3.^54^ was used to perform DSSP analysis and to extract atom coordinates from the MD trajectories. These extracted coordinates were used for calculations of interatomic distances of interest (see SI 2.4.2). For further details refer to the supplementary material.

### 3.2 Experimental methods

#### Protein preparation

Cloning of the fusion construct of h-FBP21 tWW was achieved as described in 30. For the constructs of the single h-FBP21 WW domain 1 and 2 as well as the h-FBP21 tWW-R6A mutant, please refer to SI 2.5. Protein expression was induced with IPTG at OD≈0.6 and incubated overnight at 18 °C. Protein purification was performed via affinity chromatography using HisTrap HP or GSTrap FF (GE Healthcare) based on the tag expressed in the N-terminal part of the protein, which was subsequently removed via Thrombin or TEV digestion. Further purification was achieved with size exclusion chromatography (Superdex 75 10/300 GL).

#### NMR

NMR measurements were performed on a Bruker Avance III 700 MHz spectrometer equipped with a 5 mm triple-resonance cryoprobe. ^1^H-^15^N-HSQC spectra were measured at 298 K with a protein concentration of 100 *μ*M in phosphate buffer with 10% D_2_O v/v at pH 6.7 with 8 scans and 1024 (^1^H) and 128 (^15^N) data points.

The backbone assignment for the singular WW domains of the h-FBP21 tWW was obtained from standard triple resonance spectra.

#### ITC

The Isothermal Titration Calorimetry measurements were performed at 298.15 K with the GE device MicroCaliTC200. The cell was filled up with 100 *μ*M of protein, both h-FBP21 tWW and its mutant tWW-R6A were previously dialysed in phosphate buffer. The titration was achieved with 76 injections of the proline-rich ligand of the core-splicing protein SmB/B’ (1.3 mM, Ac-GTPMGMPPPGMR-PPPPGMRGLL-NH2).

## 4 Conclusion

Using a combination of molecular simulations, peptide docking and NMR experiments we have identified and characterized two structures of the complex between the protein h-FBP21 tWW and the bivalent SmB2 ligand. The complexes, labeled complex 6 and 14, are stable over a simulation time of at least 0.5 *μs* in explicit water. In both complexes the proline-rich motifs are tightly bound to the XP grooves and exhibit the known interactions of a proline-rich sequence with an XP groove. ^1^H-^15^N-HSQC spectra showed that the residues with the largest chemical shifts were indeed those that also formed protein-ligand interactions in our complex structures.

In complex 6 the SmB2 ligand is bound in parallel orientation, while in complex 14 the ligand is bound in an antiparallel orientation. To accommodate the ligand in these two different orientations, the h-FBP21 tWW assumes two very different structures. In complex 6, the two WW domains are barely in direct contact with each other. Consequently the overall complex is rather flexible. By contrast, complex 14 forms an extensive hydrophobic interface between WW1 and WW2, and it is much more rigid than complex 6. These two different arrangements of the WW domains are made possible by the unstructured and extremely flexible linker. Our ITC experiments show that the binding of the peptide is enthalpy-driven, with entropy decreasing upon binding. Besides the loss of translational and conformational entropy for the ligand, the rigidification of linker upon binding likely contributes to the observed decrease in entropy.

We found that Arg6 in the N-terminal tail of the h-FBP21 tWW forms strikingly characteristic hydrogen bonds and ionic bridges in the cluster structures of apo-h-FBP21 tWW, as well as in our two complex structures. However, ITC and NMR experiments showed that the Arg6 does not contribute to the thermodynamic stability of the h-FBP21 tWW / SmB2 complex. These two findings are currently hard to reconcile with each other. One speculation is that the role of Arg6 is to stabilize the binding competent structure in the absence of the ligand or to stabilize an encounter complex, thereby lowering the free-energy barrier for the complex formation. In this scenario, Arg6 would influence the binding kinetics rather then the stability of the complex. In future work, this idea could be explored by Markov models^55,56^ of the binding equilibrium.

Arginine residues flanking the proline-rich motif are known to increase the specificity of ligand binding to WW domains. ^16^ In the SmB2 ligand, the C-terminal proline-rich motif is flanked by two arginines, Arg9 and Arg16. In both complexes, Arg16 forms very stable ionic bridges to glutamic acids in the h-FBP21 tWW. Mutating the arginine residues in silico, and thereby eliminating the ionic bridges, did not have an immediate effect on the stability of the two complexes.

We now have mounting evidence that h-FPB21 tWW can indeed bind the SmB2 ligand in two different orientations: parallel and antiparallel. From previous studies, we know that a spin-label at the N-terminus of the ligand interacts with both WW domains.^30^ In the present study, the 90 HADDOCK results did not show a preference for either of the ligand orientations. And finally, our workflow, which does not take the ligand orientation into account at any point, yielded two complexes which differ in the ligand orientation. This means that not only single WW domains can bind proline-rich sequences in both orientations, but this promiscuity extends to tandem WW domains. The equilibrium between parallel and antiparallel orientation might be governed by the interactions of the WW domains with residues that flank the actual proline-rich motif, such as the Arg-Glu ionic bridges in our complexes.

h-FBP21 tWW is a multivalent binder, i.e. it binds peptides with two proline-rich motifs with much higher affinity than monovalent ligands, exceeding the summed affinities of the single domains.^30,31^ Since the two WW domains are folded in the absence of a ligand and can move freely with respect to each other, this suggests an avidity-based binding mechanism. That is, binding to one WW domain increases the likelihood of binding to the second domain, because the second WW domain is tethered to the initial complex and effectively has a high local concentration. A wide variety of other binding kinetics have been reported for other tWW, including folding upon binding,^25^ and negative binding cooperativity.^57^ An atomistic model of the binding equilibrium in h-FBP21 tWW would be particularly interesting, because multivalency effects and the equilibrium between parallel and antiparallel orientation might occur on similar timescales and influence each other. Unbiased simulations of the binding kinetics are beyond the reach of current computers. However, enhanced sampling in combination with dynamical reweighting^58,59^ may open a way to a model of the binding process.

We believe that an important outcome of this study is our workflow, which consists of the following steps:

1. Decide on a list of criteria that allows you to decide whether a given complex structure is consistent with known structural characteristics of the protein-peptide complex
2. Sample the conformational ensemble of the apo-protein and classify the apo-structures using the density-based cluster algorithm CommonNN.^42,44^ (This yielded 45 apo-structures.)
3. Dock the peptide ligand to each of the apo-structures using the HADDOCK server^46–48^ (This yielded 90 proposal structures for the complex.)
4. Discard proposal structures that violate the criteria for the complex structure. Briefly simulate the other complexes to equilibrate and check stability. (In our study these were 13 complexes.)
5. Discard complexes that violate the criteria after the short simulation. The remaining structures are the results of this workflow and represent possible complex structures. They can be studied further by computational and experimental methods. (In our study, these were complexes 6 and 14.)

This workflow (Fig. 11) can be applied to any binding process that occurs predominantly via conformational selection. This restriction arises, because the protein-protein docking in step 3 essentially assumes a conformational selection mechanism. In our workflow, induced fit effects in the binding process are partially accounted for by an extensive sampling of the apo-structure in step 2 and by simulating the proposed protein-protein complex structures in step 4, which allows the complex to re-adjust if necessary. Additionally, the docking protocol itself can account for induced fit effects to some extent by allowing for some flexibility in the two binding partners or by using experimental interaction data to drive the docking^46,60^ In step 3, HADDOCK^46,60^ can in principle be replaced by other protein-protein docking or peptide docking methods. According to recent results from CAPRI (Critical Assessment of Predicted Interactions), a well-established community-wide experiment to benchmark docking methods, HADDOCK ranks among the the best-performing docking methods. ^61^ Close contesters are ClusPro,^62^ HDock,^63^ MDOCKPP,^64^ and LZerd.^65^ Each of these docking methods expects a globular protein as the receptor protein. By embedding the docking step in our workflow, one can predict protein-protein complexes in which the receptor is either very flexible or consists of sub-domains that can assume several relative orientations. The workflow is particularly useful to elucidate complex structures of repeats of protein domains, because the apo-conformations of these proteins can be characterized in terms of the geometric coordinates that describe the relative position and orientation of these domains. The workflow allows us to systematically generate possible complex structures for repeats of recognition domains, such as WW domains and SH3 domains. These structures will help us understand how interactions between the domains, interactions between the domains and the ligand binding motifs, as well as synergistic and multivalent effects shape the astonishing versatility and specifcity of protein-protein interactions.

**Figure 11:**
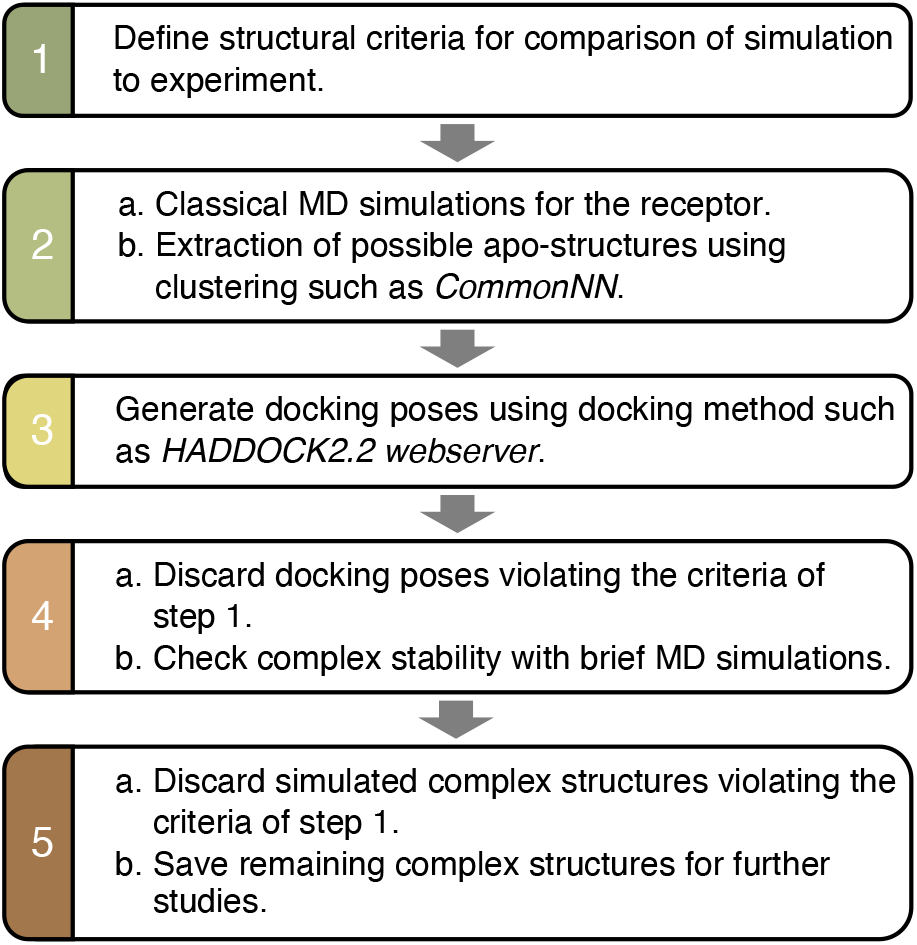
Overview of our approach on the identification of complex structures for the h-FBP21 tWW with a proline-rich ligand from SmB/B’ core-splicing protein.

## 5 Data and software availability

The MD data including topology files, trajectories, clustering results and docking structures are available from the corresponding author (B.G.K.) upon reasonable request. Structures for the h-FBP21 tWW / SmB2 complexes 6 and 14 are available as pdf-and pdb-files at DOI (doi:10.5281/zenodo.5680225). Software used within this study is indicated in section 3. Further details regarding parameters, settings and their use for the analyses performed in this study are provided in the SI. Requests for custom scripts should be directed to the corresponding author (B.G.K.). The free software used within this study is available under the following links: GROMACS (https://manual.gromacs.org/documentation/), Python (https://www.python.org/downloads), CommonNN algorithm (https://github.com/janjoswig/CommonNNClustering), HADDOCK2.2 webserver (https://alcazar.science.uu.nl/services/HADDOCK2.2/), Avogadro (https://avogadro.cc), VMD (https://www.ks.uiuc.edu/Research/vmd/). The NMR spectra were processed by the free software CcpNmr Analysis (https://www.ccpn.ac.uk/v2-software/software/analysis) and the commercial software TopSpin by Bruker. For the analysis of ITC data, the commercial software MicroCal by Malvern Panalytical was used.

## Supporting information

The SI is freely available in pdf format together with this manuscript. It includes additional data and detailed descriptions of all computational and experimental methods used in this study. The SI also provides explanations for decisions made during the analyses.

## Acknowledgement

This work was funded by Deutsche Forschungsgemeinschaft (DFG - German Research Foundation): project ID 392923329 (RTG 2473), project ID 387284271 (SFB 1349) and project ID 32049920 (SFB 765).

The authors thank the Zentraleinrichtung für Datenverarbeitung (ZEDAT) of Freie Universitat Berlin for computing time. For NMR experiments, the authors acknowledge the assistance of the Core Facility BioSupramol supported by the DFG.

## Advancing women in science

The authors believe that the current challenges, such as climate change, scarcity of energy and resources, and global health hazards, can only be addressed if the most capable persons from all socio-economic and cultural backgrounds have equal access to STEM education and an equal opportunity to pursuit a career in these fields. We are committed to promote diversity, gender equality and to create a family-friendly environment in our labs. Our efforts are embedded in long-standing and successful diversity and gender equality policies at the Freie Universität Berlin, for which the university has been recognized with several awards.

## Notes

### Competing Interest Statement

The authors have declared no competing interest.

### Summary of Updates

Final version for publication.

