## Supplemental Information for "Target Recognition in Tandem WW Domains: Complex Structures for Parallel and Antiparallel Ligand Orientation in h-FBP21 Tandem WW"

### 1 Additional data

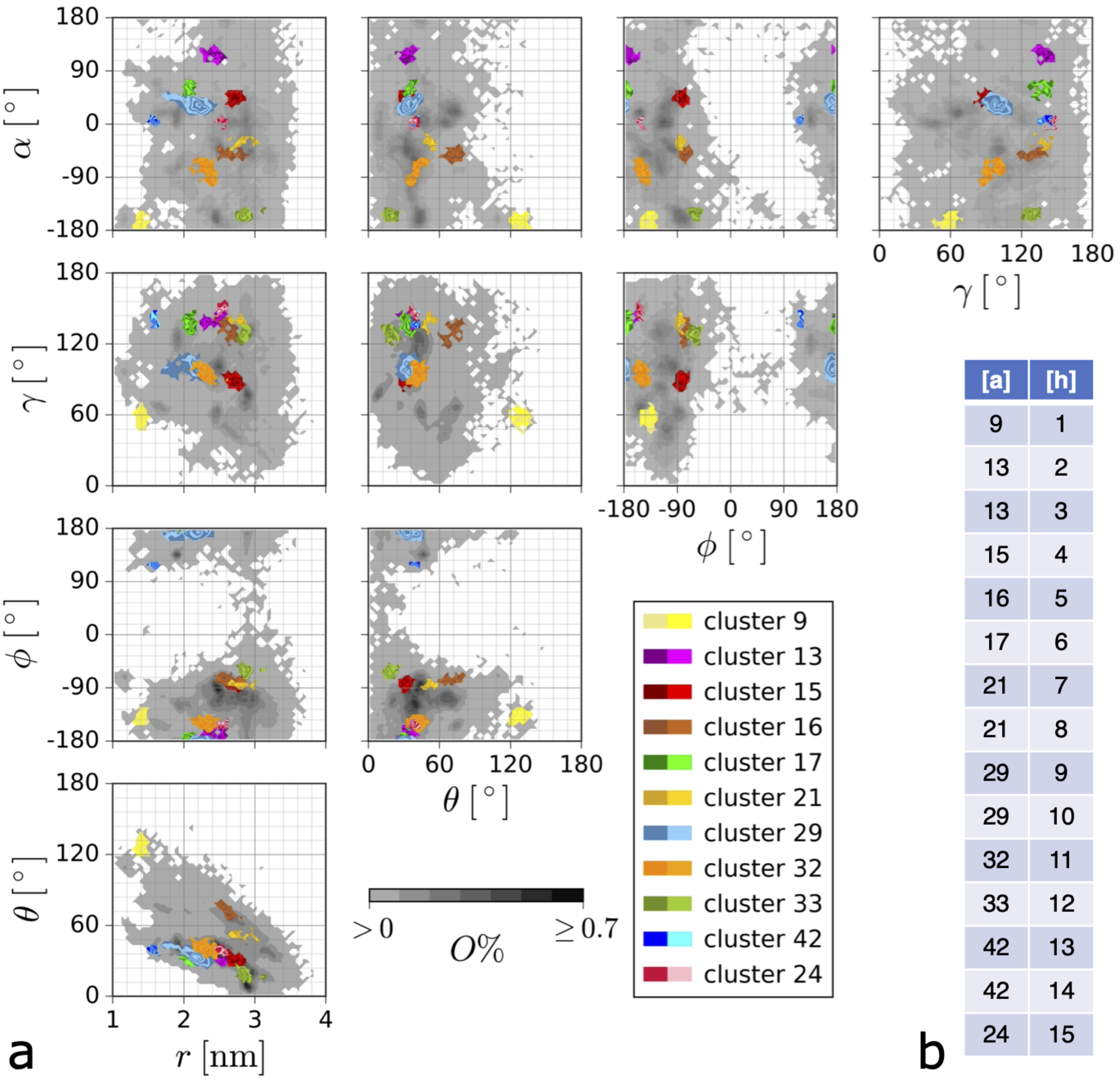


| [a] | [h] |
|---|---|
| 9 | 1 |
| 13 | 2 |
| 13 | 3 |
| 15 | 4 |
| 16 | 5 |
| 17 | 6 |
| 21 | 7 |
| 21 | 8 |
| 29 | 9 |
| 29 | 10 |
| 32 | 11 |
| 33 | 12 |
| 42 | 13 |
| 42 | 14 |
| 24 | 15 |

Figure S1: (a) Clusters obtained with CommonNN clustering on the MD data set of the apo-h-FBP21 tWW domain. Clusters 1-15 (corresponding to Table 1 and Fig. 5 in the main text). Clusters are projected into pairs of the reaction coordinates $(r, \theta, \phi), \alpha, \gamma$. (b) This panel (b) shows the map between cluster number [a] and complex structure number [h].

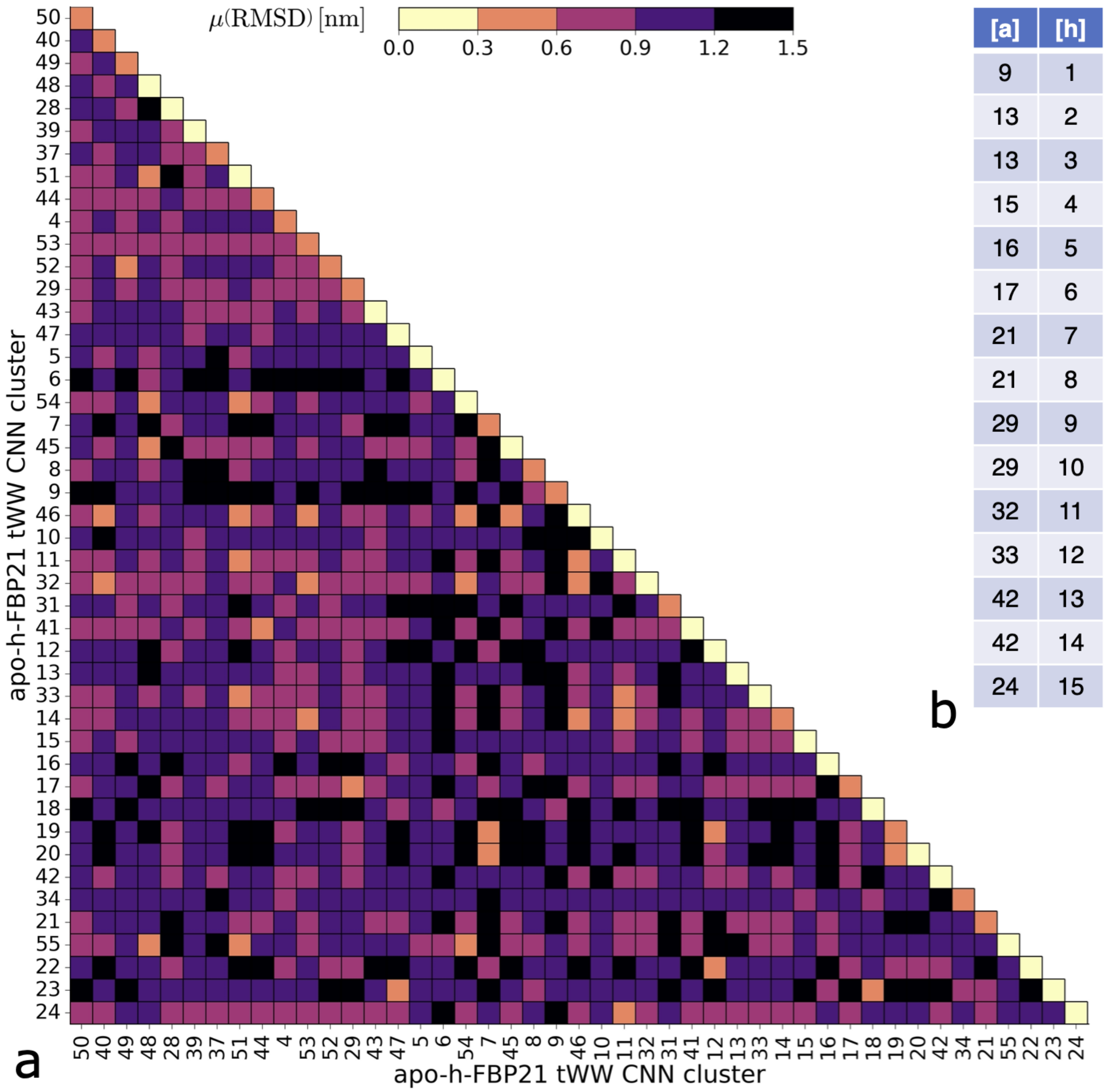


| [a] | [h] |
|---|---|
| 9 | 1 |
| 13 | 2 |
| 13 | 3 |
| 15 | 4 |
| 16 | 5 |
| 17 | 6 |
| 21 | 7 |
| 21 | 8 |
| 29 | 9 |
| 29 | 10 |
| 32 | 11 |
| 33 | 12 |
| 42 | 13 |
| 42 | 14 |
| 24 | 15 |

Figure S2: (a) Structural difference between the clusters of apo-h-FBP21 tWW. Each element $m_{ij}$ shows the average pairwise $C_\alpha$-RMSD between structures form cluster $i$ and $j$. The clusters are sorted by size in decreasing order: top to bottom, left to right. Panel (a) uses the cluster numbers from the CommonNN analysis. After the HADDOCK docking, we numbered the selected complex structures 1 to 15. This panel (b) shows the map between cluster number [a] and complex structure number [h].

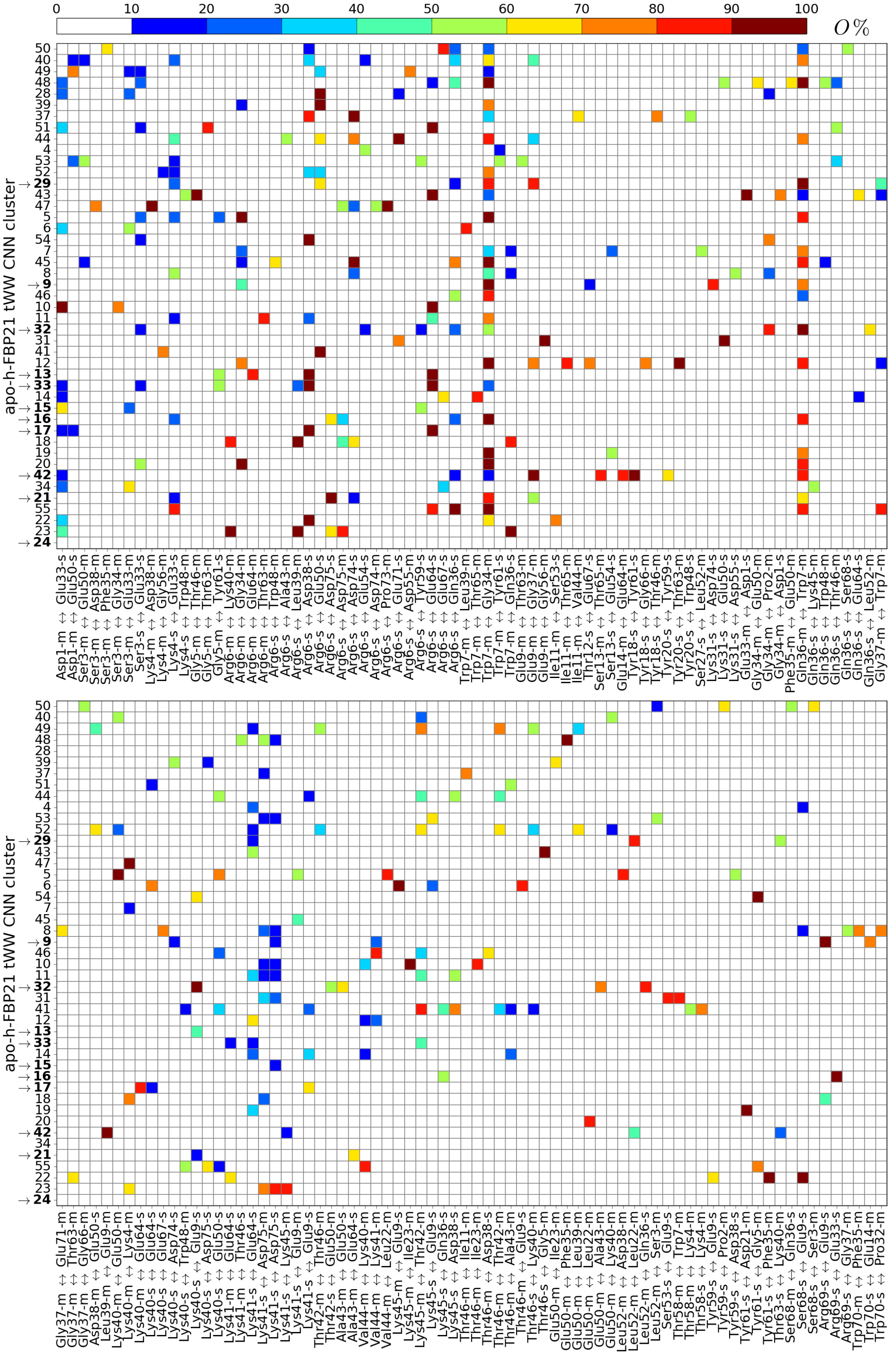


Figure S3: Interdomain hydrogen bonds the clusters of apo-h-FBP21 tWW. "donor↔acceptor", with a relative occurrence of > 50% in at least one of the h-FBP21 tWW CommonNN clusters indicated on the ordinate. All hydrogen bonds are classified as main chain ('m') or side chain ('s') interactions. Clusters, which were used for MD simulations of tWW+SmB2 complexes are highlighted in bold.

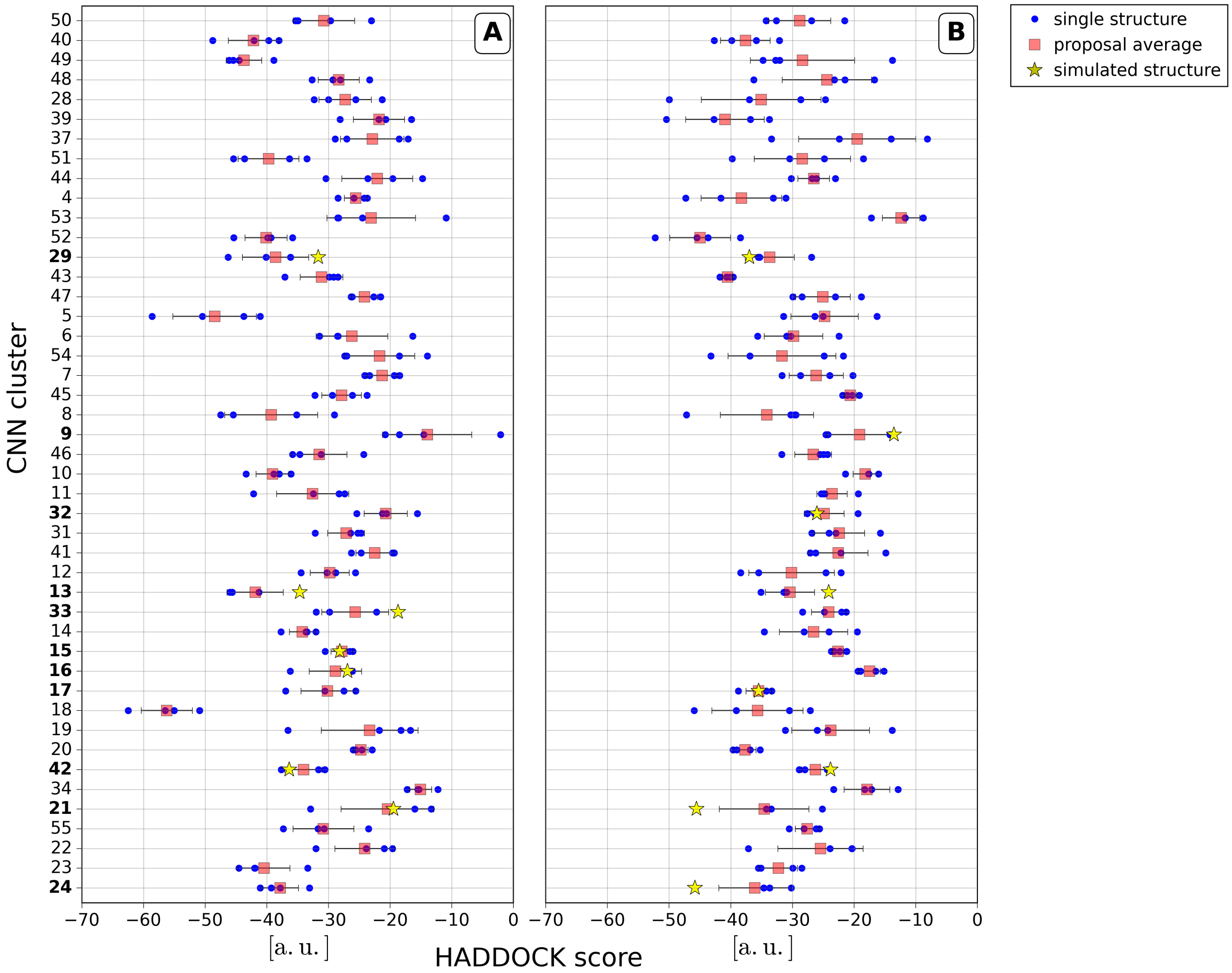


Figure S4: HADDOCK docking results between the SMB2 ligand and structures drawn form the 45 CommonNN clusters of apo-h-FBP21 tWW. (A) HADDOCK proposal with lowest HADDOCK score, (B) HADDOCK proposal with second lowest HADDOCK score The results are ordered with respect to the size of the cluster from which the apo-structure was drawn (decreasing size from top to bottom). Each HADDOCK proposal contains four complex structures. Blue dots: HADDOCK scores of the four complex structures within one proposal. Red square: average HADDOCK score of the four complex structures. The corresponding CommonNN cluster number is printed in bold. Yellow star: complex structures used for further analysis.

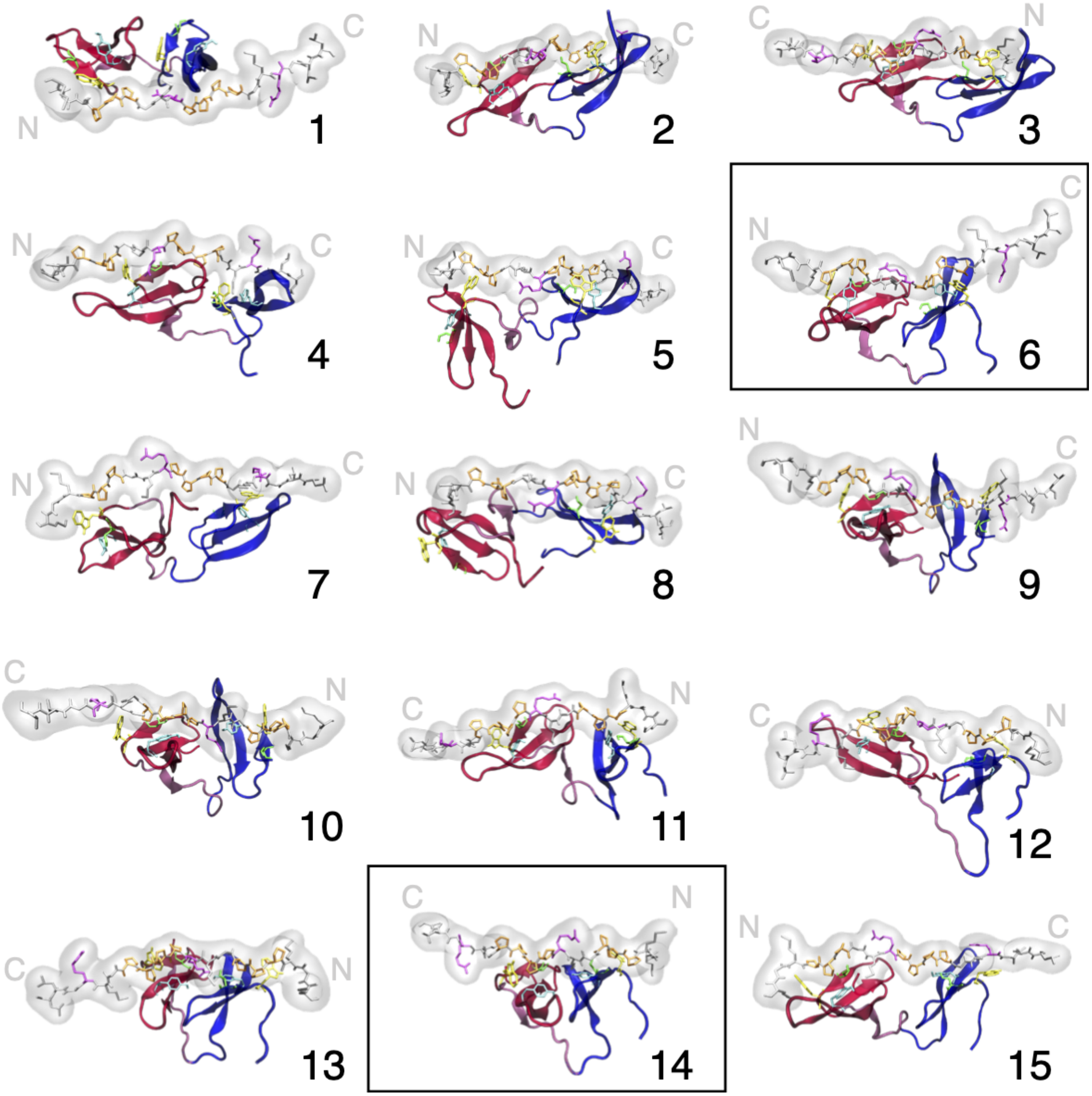


Figure S5: Selected structures for a complex of h-FBP21 tWW domain and SmB2 ligand, that were obtained by docking with HADDOCK2.2 webserver and subsequently simulated over 50 ns. The h-FBP21 tWW domain is coloured according to its segments and residues relevant for binding (XP grooves) are pictured: WW domain 1 (red), interdomain region (pink), WW domain 2 (blue), XP grooves: Trp (yellow), Ser (green), Tyr (cyan) The SmB2 ligand is coloured in gray, residues belonging to the proline-rich motifs are coloured according to residue type: Pro (orange), Arg (magenta). The C- and N-terminus of the SmB2 ligand are labelled for each complex structure. The two best complex structures are highlighted in the black rectangles.

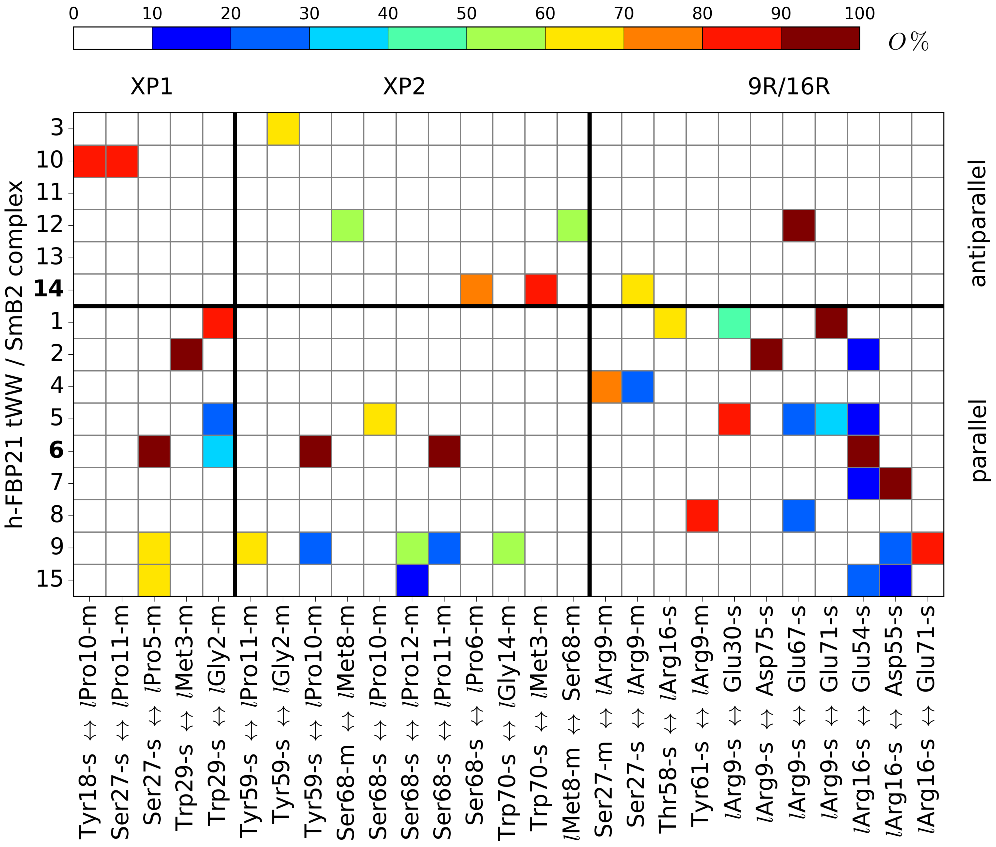


Figure S6: Intermolecular hydrogen bonds denoted as "donor↔acceptor", with a relative occurrence of > 50% in at least one of the simulated HADDOCK proposals indicated on the ordinate. All hydrogen bonds are classified as main chain ('m') or side chain ('s') interactions. '*l*' denotes residues belonging to the SmB2 ligand. The complexes are differentiated based on the orientation of the SmB2 ligand: 'parallel', 'antiparallel'. All hydrogen bonds are sorted in three groups: XP1: hydrogen bonds involving the residues Tyr18, Ser27 or Trp29; XP2: hydrogen bonds involving the residues Tyr59, Ser68 or Trp29; 9R/16R: hydrogen bonds involving the residues Arg9 and Arg16 of the SmB2 ligand. Hydrogen bonds involving both, members of the XP grooves and Arg9/16 of the SmB2 ligand, are solely presented in '9R/16R'.

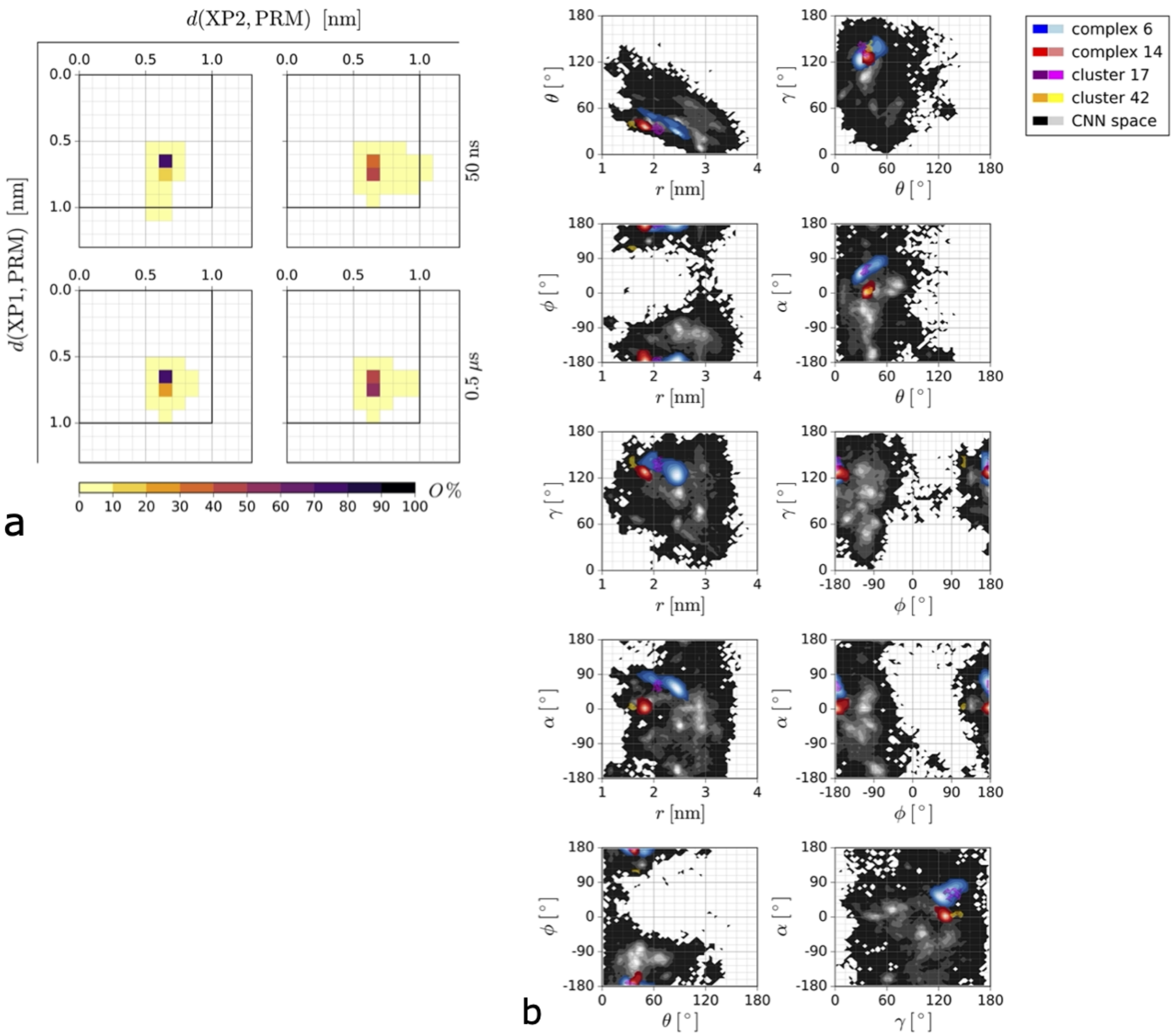


Figure S7: (a) Joint distributions for the distances between the XP grooves of WW domain 1 (XP1) and WW domain 2 (XP2) to the respective closest proline-rich motif (PRM). (b) Distribution of the tWW structures in the complexes 6, 14 and the CommonNN clusters 17 and 42 projected onto pairs of reaction coordinates $(r, \theta, \phi), \alpha, \gamma$.

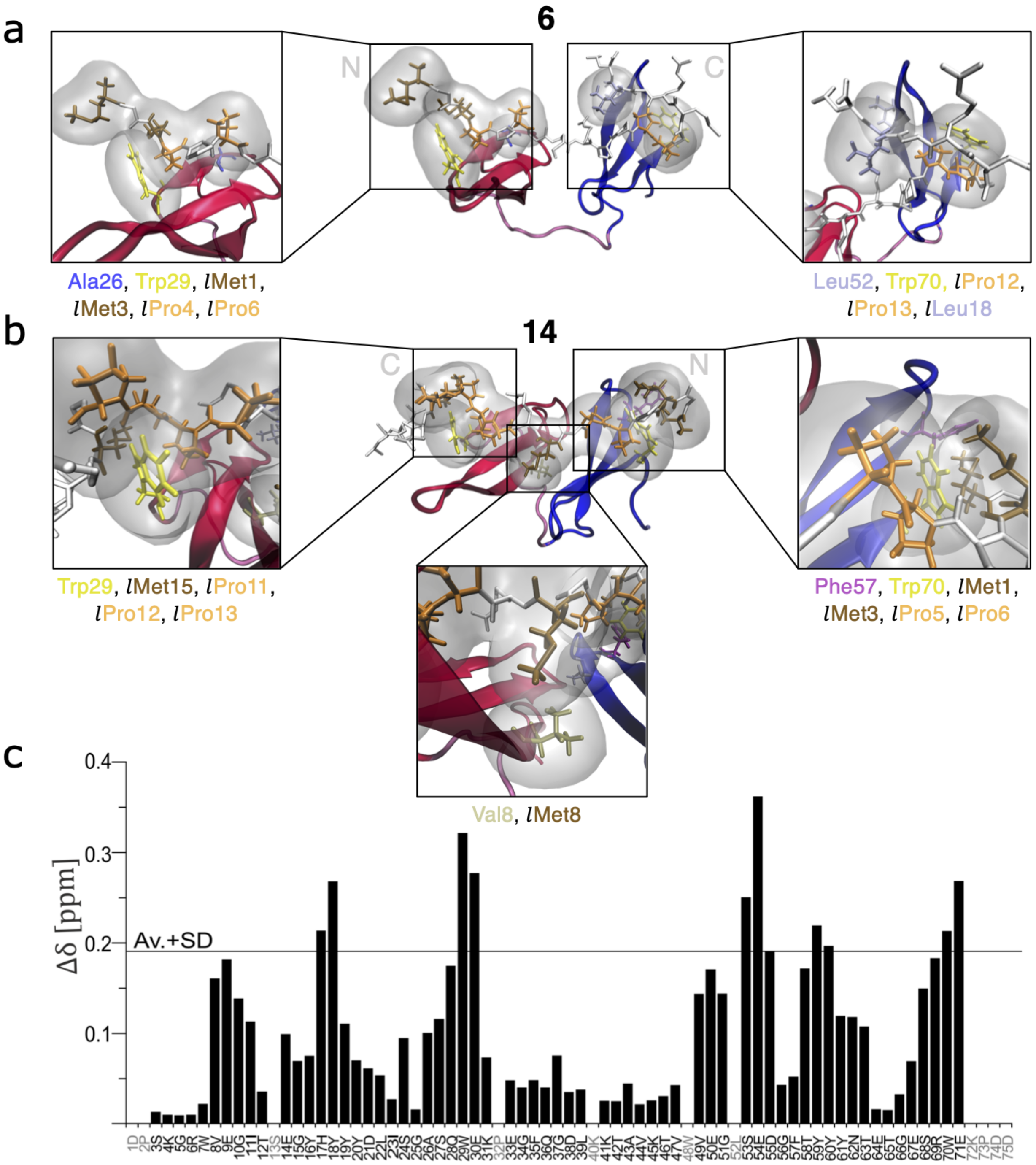


Figure S8: (a,b) Hydrophobic contacts in the h-FBP21 tWW / SmB2 complex structures 6 (a) and 14 (b). Residues considered significantly involved in the hydrophobic contacts are highlighted according to residue type: blue: alanine, yellow: tryptophan, brown: methionine, orange: proline, pale-blue: leucine, ochre: valine, purple: phenylanaline. (c) Chemical shift differences in $^{1}H$-$^{15}N$-HSQC spectra of ligand-free (apo) and ligand-bound (holo) h-FBP21 tWW. Unassigned residues are colored in grey. Chemical shift differences above the average plus standard deviation (solid black line) are considered significant.

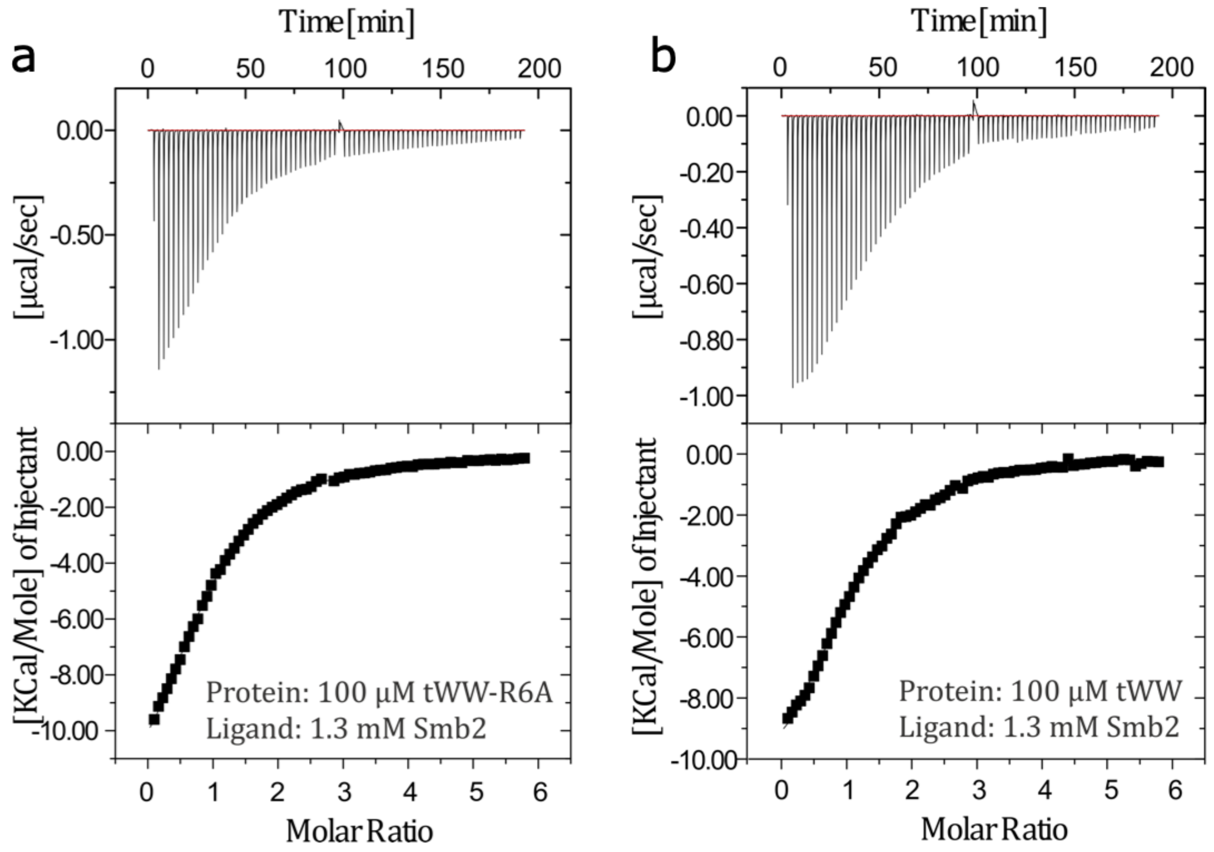


Figure S9: An Isothermal Titration Calorimetry (ITC) assay to detect the thermodynamic parameter of the interaction between the SmB2 ligand and the wild type h-FBP21 tWW or its mutated form tWW-R6A.

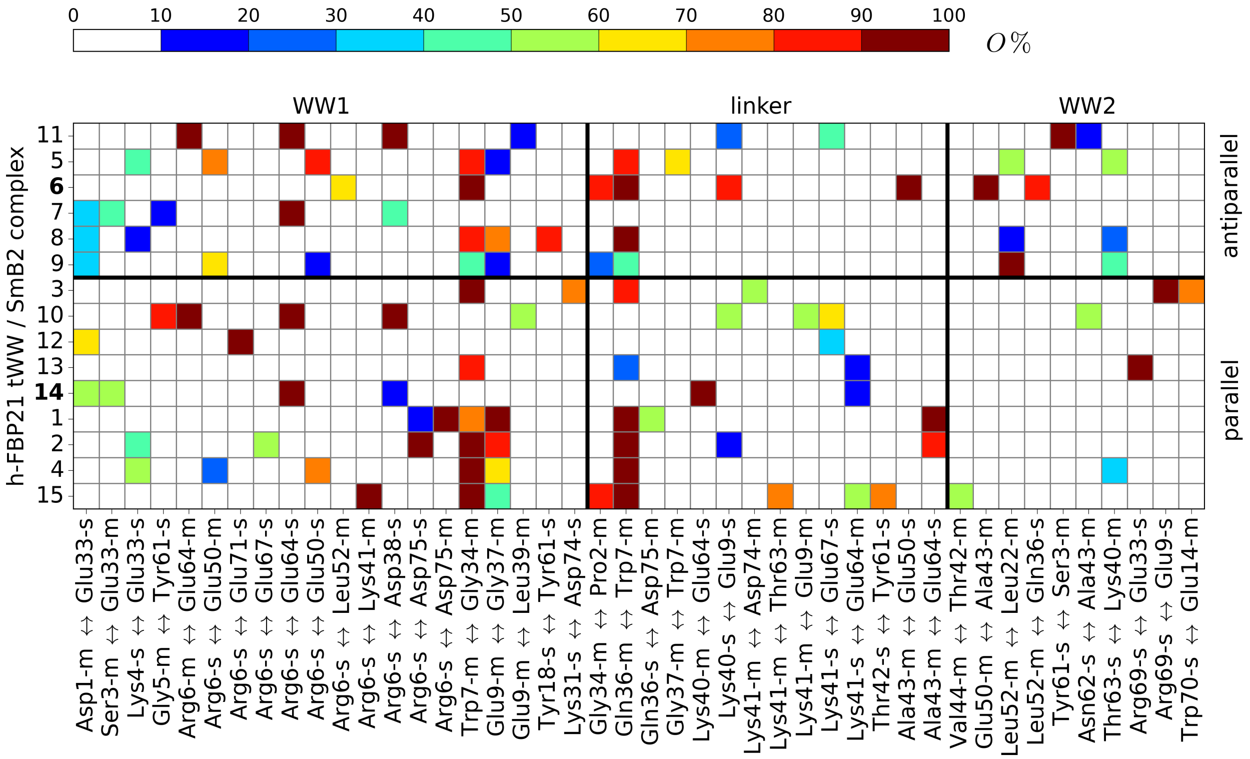


Figure S10: Interdomain hydrogen bonds denoted as "donor↔acceptor", with a relative occurrence of $> 50\%$ in at least one of the simulated HADDOCK proposals indicated on the ordinate. The complexes are differentiated based on the orientation of the SmB2 ligand: 'parallel', 'antiparallel'. The hydrogen bonds are sorted into three groups in accordance to the belonging of the donor to either WW domain 1 (WW1), interdomain region (linker) and WW domain 2 (WW2).

Table S1: List of resonances giving rise to NOE crosspeaks between amide groups in a $^{1}H$-$^{15}N$-NOESY-HSQC spectrum of h-FBP21 tWW in the presence of SmB2 ligand. The distance estimated from NOE crosspeak intensities is given for amide proton pairs.

| Resonances | estimated distance [Å] | upper limit [Å] | lower limit [Å] | NOE crosspeak intensity [a.u.] |
|---|---|---|---|---|
| 4LysH32SerH | 3.61784 | 4.34141 | 2.89427 | 1.10E+06 |
| 4LysH-3SerH | 3.77842 | 4.5341 | 3.02274 | 8.47E+05 |
| 11IleH-12ThrH | 4.21707 | 5.06048 | 3.37365 | 4.38E+05 |
| 11IleH-12ThrH | 4.05524 | 4.86629 | 3.24419 | 5.54E+05 |
| 12ThrH-16TyrH | 3.70219 | 4.44263 | 2.96175 | 9.58E+05 |
| 12ThrH-16TyrH | 3.51638 | 4.21965 | 2.8131 | 1.30E+06 |
| 15GlyH-16TyrH | 3.08583 | 3.70299 | 2.46866 | 2.86E+06 |
| 15GlyH-16TyrH | 3.01707 | 3.62048 | 2.41365 | 3.27E+06 |
| 23IleH-22LeuH | 3.47492 | 4.1699 | 2.77993 | 1.40E+06 |
| 23IleH-22LeuH | 3.51095 | 4.21314 | 2.80876 | 1.32E+06 |
| 24SerH-23IleH | 3.14574 | 3.77489 | 2.51659 | 2.54E+06 |
| 24SerH-23IleH | 3.14136 | 3.76963 | 2.51309 | 2.57E+06 |
| 25GlyH-24SerH | 3.1536 | 3.78432 | 2.52288 | 2.51E+06 |
| 25GlyH-24SerH | 3.15864 | 3.79036 | 2.52691 | 2.48E+06 |
| 25GlyH-26AlaH | 3.2771 | 3.93253 | 2.62168 | 1.99E+06 |
| 25GlyH-26AlaH | 3.03745 | 3.64494 | 2.42996 | 3.14E+06 |
| 28GlnH-19TyrH | 3.90051 | 4.68061 | 3.12041 | 7.00E+05 |
| 28GlnH-19TyrH | 3.62833 | 4.35399 | 2.90266 | 1.08E+06 |
| 29TrpH-30GluH | 3.37387 | 4.04864 | 2.69909 | 1.67E+06 |
| 29TrpH-30GluH | 3.09919 | 3.71902 | 2.47935 | 2.78E+06 |
| 35PheH-34GlyH | 3.7168 | 4.46016 | 2.97344 | 9.35E+05 |
| 35PheH-34GlyH | 4.24945 | 5.09933 | 3.39956 | 4.19E+05 |
| 42ThrH-43AlaH | 3.84446 | 4.61335 | 3.07557 | 7.64E+05 |
| 42ThrH-43AlaH | 3.86061 | 4.63273 | 3.08848 | 7.45E+05 |
| 44ValH-45LysH | 3.88341 | 4.66009 | 3.10673 | 7.19E+05 |
| 44ValH-45LysH | 3.89037 | 4.66844 | 3.1123 | 7.11E+05 |
| 53SerH-57PheH | 3.51496 | 4.21795 | 2.81197 | 1.31E+06 |
| 53SerH-57PheH | 3.50739 | 4.20887 | 2.80591 | 1.32E+06 |
| 55AspH-54GluH | 4.27776 | 5.13331 | 3.42221 | 4.02E+05 |
| 55AspH-54GluH | 4.12858 | 4.9543 | 3.30287 | 4.98E+05 |
| 55AspH-56GlyH | 3.41517 | 4.0982 | 2.73214 | 1.55E+06 |
| 55AspH-56GlyH | 3.08475 | 3.7017 | 2.4678 | 2.86E+06 |
| 63ThrH-49ValH | 4.57757 | 5.49308 | 3.66206 | 2.68E+05 |
| 63ThrH-49ValH | 4.45492 | 5.3459 | 3.56393 | 3.15E+05 |

Table S1 (continuation): List of resonances giving rise to NOE crosspeaks between amide groups in a $^{1}H$-$^{15}N$-NOESY-HSQC spectrum of h-FBP21 tWW in the presence of SmB2 ligand. The distance estimated from NOE crosspeak intensities is given for amide proton pairs.

| Resonances | estimated distance [Å] | upper limit [Å] | lower limit [Å] | NOE crosspeak intensity [a.u.] |
|---|---|---|---|---|
| 64GluH-65ThrH | 2.87523 | 3.45028 | 2.30018 | 4.36E+06 |
| 64GluH-65ThrH | 2.8227 | 3.38724 | 2.25816 | 4.88E+06 |
| 65ThrH-67GluH | 3.82704 | 4.59245 | 3.06163 | 7.85E+05 |
| 66GlyH-65ThrH | 2.93859 | 3.52631 | 2.35087 | 3.83E+06 |
| 66GlyH-65ThrH | 2.90793 | 3.48952 | 2.32634 | 4.08E+06 |
| 66GlyH-67GluH | 3.26378 | 3.91654 | 2.61103 | 2.04E+06 |
| 66GlyH-67GluH | 3.29999 | 3.95999 | 2.63999 | 1.91E+06 |
| 69ArgH-60TyrH | 3.8163 | 4.57956 | 3.05304 | 7.98E+05 |
| 69ArgH-60TyrH | 3.69768 | 4.43722 | 2.95814 | 9.65E+05 |
| 71GluH-70TrpH | 3.22774 | 3.87329 | 2.58219 | 2.18E+06 |
| 71GluH-70TrpH | 3.10075 | 3.7209 | 2.4806 | 2.77E+06 |

### 2 Methods

#### 2.1 MD simulations

We conducted classical, all-atom molecular-dynamics simulations of the apo-h-FBP21 tWW (see 2.1.1), and of the h-FBP21 tWW in complex with the (mutated) SmB2 peptide (see 2.1.2 and 2.1.3). See Table S2 for a summary of the simulations.

##### 2.1.1 apo-h-FBP21 tWW

MD simulations of the apo-h-FBP21 tWW were performed with GROMACS 5.0.2 simulation package[1–3] in explicit water. The protein was modelled using the AMBER ff99SB*-ILDNP force field.[4] The starting structure (PDB entry: 2JXW[5]) was first energy minimised in vacuum using the steepest descent algorithm[1] (emtol=1000 kJ/(mol· nm), nsteps=5000). It was then solvated in TIP3P[6] water in a dodecahedronal box ($V$=291.64 nm$^3$). 7 $Na^+$ ions were added to obtain a box with neutral electric charge. The solvated system was energy minimised (steepest-descent), followed by $NVT$ and $NpT$ equilibrations for 100 ps at 300 K (d$t$=2 fs) with periodic boundary conditions in all directions. For data production, the system was propagated using leap-frog integration[7] with an integration time step d$t$=2 fs. All bonds were constrained using the LINCS algorithm[8] (iter=1,order=4). Periodic boundary conditions were applied in all three directions. The simulations were conducted in the $NpT$ ensemble at temperature $T$=300 K (velocity-rescale thermostat,[9] coupling time $\tau_T$=0.1 ps) and $p$=1 bar (Parrinello-Rahman barostat,[10] coupling time $\tau_p$=2 ps). Van-der-Waals interactions were limited with Verlet cut-off scheme[11] with a cut-off radius $r_{\mathrm{vdw}}$= 1 nm. The neighbourlist was updated every nstlist=5 integration time steps. The Coulomb interactions were calculated using the Particle-Mesh-Ewald algorithm[12] (pme-order=4, fourierspacing=0.16 nm, cut-off for the short-range electrostatic interactions $r_{\mathrm{coulomb}}$=1 nm. Solute coordinates were written to file every 1 ps. 70 trajectories with an average simulation length of 0.5±0.2 $\mu$s were produced yielding a total simulation time of 35.05 $\mu$s.

##### 2.1.2 HADDOCK h-FBP21 tWW / SmB2 complexes

Out of the 360 h-FBP21 tWW / SmB2 complex structures obtained with HADDOCK2.2 webserver[13,14] (see 2.3), 15 were selected as starting structures for MD simulations with GROMACS 2016.5[2,3,15] (for a visualisation of these starting structures please refer to Fig. S5). All complex structures were modelled using the AMBER ff99SB*-ILDNP force field.[4] They were energy minimised in vacuum with steepest descent algorithm[15] (emtol=1000 kJ/(mol·nm), nsteps=50000). Then, the complex structures were solvated in TIP3P[6] water in dodecahedronal boxes with average volume $V_{\mathrm{mean}}$=456.00±51.96 nm$^3$. 5 $Na^+$ ions were added to ensure charge balance. The solvated complexes were energy minimised (steepest descent), followed by $NVT$ and $NpT$ equilibrations for 150 ps at 300 K (d$t$=2 fs) with periodic boundary conditions in all directions. The parameters of the production runs were as in 2.1.1 except for the neighbour list updating frequency nstlist=10 (Verlet-cut-off scheme).

For each starting structure, 50 ns of simulation data were generated saving the solute coordinates every 1 ps.

##### 2.1.3 h-FBP21 tWW / SmB2 complexes (6, 14) and mutation studies

**Complexes 6 and 14 with wild-type ligand (SmB2-WT).** For the h-FBP21 tWW / SmB2 complexes 6 and 14 with SmB2-WT as ligand, MD simulations were conducted with GROMACS 2016.5[2,3,15] and GROMACS 2019 release version.[16] Starting structures that fulfilled the distance criterion explained in 2.4.3, were extracted from the preceding MD simulations of the HADDOCK h-FBP21 tWW / SmB2 complexes described in 2.1.2. Each starting structure was modelled using the AMBER ff99SB*-ILDNP force field.[4] The system setup, energy minimisation and solvation in TIP3P[6] followed the protocol described in 2.1.2. $NVT$ and $NpT$ equilibrations were carried out for 100 ps at 300 K (d$t$=2 fs) with periodic boundary conditions in all directions. In the production MD simulations, the system was propagated using leap-frog integration[7] with an integration time step d$t$=2 fs. Bonds including hydrogen atoms were constrained using the LINCS algorithm[8] (iter=1, order=4).

The simulations were first conducted in the $NpT$ ensemble at temperature $T$ = 300 K (velocity-rescale thermostat,[9] coupling time $\tau_T$=0.01 ps) and $p$ = 1 bar (Parrinello-Rahman barostat,[10] coupling time $\tau_p$=2 ps) for 0.25 $\mu$s. Then, the simulations were continued in the $NVT$ ensemble at temperature $T$ = 300 K (velocity-rescale thermostat,[9] coupling time $\tau_T$=0.01 ps) for 0.25 $\mu$s. In both ensembles, Van-der-Waals interactions were truncated using the Verlet cut-off scheme[11] with a cut-off radius of $r_{\text{vdw}}$ = 1 nm. The neighbourlist was updated every nstlist=10 integration timesteps. The Coulomb interactions were calculated using the Particle-Mesh-Ewald algorithm[12] (pme-order=4, fourierspacing=0.16 nm, cut-off for the short-range electrostatic interactions $r_{\text{coulomb}}$=1 nm). Solute coordinates were written to file every 1 ps. In total, 0.5 $\mu$s of simulation data were generated for each starting structure (Table S2).

**Complexes 6 and 14 with mutated ligand.** In the starting structures for the complexes 6 and 14 (see 2.1.3), the following amino acids were replaced in the SmB2 ligand:

- Arg9 by Ala9 (SmB2-R9A),
- Arg16 by Ala16 (SmB2-R16A),
- Arg9 by Ala9 and Arg16 by Ala16 (SmB2-R9A/R16A).

All modifications were made with the chemical editor AVOGADRO.[17] The resulting 12 starting structures were simulated for 0.5 $\mu$s (Table S2) according to the procedure described for the wild type complexes (see 2.1.3).

Table S2: Overview of the performed MD simulations with temperature $T$, average volume $V$ and simulation times $t$.

| system | specification | $T$ [K] | $V$ [nm$^3$] | $t$ [$\mu$s] |
|---|---|---|---|---|
| apo-h-FBP21 tWW | PDB entry: 2JXW[5] | 300 | 291.64 | 35.05 |
| h-FBP21 tWW / SmB2 complex | structures from HADDOCK | 300 | 456 | 0.05 |
| h-FBP21 tWW / SmB2 complex (6) | SmB2-WT | 300 | 357.12 | 0.5 |
| | SmB2-R9A | 300 | 358.86 | 0.5 |
| | SmB2-R16A | 300 | 250.84 | 0.5 |
| | SmB2-R9A/R16A | 300 | 250.88 | 0.5 |
| h-FBP21 tWW / SmB2 complex (14) | SmB2-WT | 300 | 316.99 | 0.5 |
| | SmB2-R9A | 300 | 317.08 | 0.5 |
| | SmB2-R16A | 300 | 222.50 | 0.5 |
| | SmB2-R9A/R16A | 300 | 222.42 | 0.5 |

#### 2.2 Analysis of the apo-h-FBP21 tWW simulations

In preparation for the analyses, all trajectories were modified with GROMACS so that the h-FBP21 tWW was centered in the box (gmx trjconv -pbc mol -center). An additional rotational and translational fit was applied including a least-squares fit on the C_$\alpha$-atoms (gmx trjconv -fit rot+trans).[16]

##### 2.2.1 DSSP plots

The temporal development of the secondary structure of the apo-h-FBP21 tWW was examined with the feature `mdtraj.compute_dssp` out of the python library `mdtraj` 1.9.3.,[18] which re-implements the DSSP 2.2.0 program.[19,20]

##### 2.2.2 Removing outliers from the simulation data set of the apo-h-FBP21 tWW

On the basis of the distances between the $C_\alpha$ atoms of the following pairs of residues: His17, Thr58 ($d_1$), His17, Tyr61 ($d_2$) and Tyr20, Tyr61 ($d_3$), the original simulation data set

($35.05 \cdot 10^6$ structures) was divided up into two data sets $C$ and $E$ defined as follows:

$$E = \{S \mid d_1 > 2.65\,\mathrm{nm} \;\&\; d_2 > 2.65\,\mathrm{nm} \;\&\; d_3 > 2.65\,\mathrm{nm}\}$$

$$C = S \backslash E.$$

$C$ comprises $31 \cdot 10^6$ structures (89.1% of the original data set) and $E$ comprises $3.81 \cdot 10^6$ structures (10.9% of the original data set).

Introducing a fourth distance between the $C_\alpha$ atoms of Tyr20,Thr58 ($d_4$), the data set $E$ was split up again as follows:

$$E_2 = \{E \mid d_1 > 2.65\,\mathrm{nm} \;\&\; d_2 > 2.65\,\mathrm{nm} \;\&\; d_3 > 2.65\,\mathrm{nm} \;\&\; d_4 > 2.65\,\mathrm{nm}\}$$

$$E_1 = E \backslash E_2,$$

where $E_2$ comprises $3.54 \cdot 10^6$ structures (10.1% of the original data set) and $E_1$ comprises $0.27 \cdot 10^6$ structures (0.7% of the original data set). The data set of compact structures, which was used for further analyses includes the data sets $C$ and $E_1$ (in total: $31.51 \cdot 10^6$ structures, 89.9%), whereas $E2$ represents the data set of extended structures.

##### 2.2.3 Translational and rotational fit - reaction coordinates

To guarantee the same spatial orientation of the $\beta$-sheet of WW domain 1 in all h-FBP21 tWW structures, the position vector $\mathbf{r}_i$ of every atom $i$ was transformed as follows:

$$\mathbf{r}_i^{\mathrm{trans,rot}} = \mathbf{R}(\alpha) \left[\mathbf{r}_i - \mathbf{r}^{\mathrm{ref}}\right], \tag{1}$$

$$\mathbf{r}_i^{\mathrm{trans,rot}} = (x_i, y_i, z_i)^{\mathrm{T}} \xrightarrow[\text{transformation}]{\text{coordinate}} \mathbf{r}_i^{\mathrm{trans,rot}} = (r, \theta, \phi)^{\mathrm{T}} \tag{2}$$

$$\mathbf{r}_i^{\mathrm{final}} = \mathbf{R}(\phi)\mathbf{r}_i^{\mathrm{trans,rot}}. \tag{3}$$

First, the translational shift $\left[\mathbf{r}_i - \mathbf{r}^{\mathrm{ref}}\right]$ with $\mathbf{r}^{\mathrm{ref}}$ being the position of the $C_\alpha$(His17) as returned by the MD simulation, shifts $C_\alpha$(His17) to the origin of the coordinate system. The rotation matrix $\mathbf{R}(\alpha)$ rotates the distance vector $\mathbf{r}_{17,20}$ between $C_\alpha$(His17) and $C_\alpha$(Tyr20) onto the positive $z$-axis and is given as

$$\mathbf{R}(\alpha) = \begin{bmatrix} n_x^2 b + \cos(\alpha) & n_x n_y b - n_z \sin(\alpha) & n_x n_z b + n_y \sin(\alpha) \\ n_y n_x b + n_z \sin(\alpha) & n_y^2 b + \cos(\alpha) & n_y n_z b - n_x \sin(\alpha) \\ n_z n_x b - n_y \sin(\alpha) & n_z n_y b + n_x \sin(\alpha) & n_z^2 b + \cos(\alpha) \end{bmatrix}, \tag{4}$$

with

$$\begin{aligned} \mathbf{n} &= \left( \frac{\mathbf{r}_{17,20} \times \mathbf{u}}{||\mathbf{r}_{17,20} \times \mathbf{u}||} \right) \\ \alpha &= \arccos \left( \frac{\mathbf{r}_{17,20} \cdot \mathbf{u}}{||\mathbf{r}_{17,20}|| \cdot ||\mathbf{u}||} \right) \\ b &= 1 - \cos(\alpha), \end{aligned} \tag{5}$$

where $\mathbf{u} = (0, 0, 1)^{\mathrm{T}}$ denotes the unit vector in positive $z$-direction.

Second, all position vectors $\mathbf{r}_i^{\text{trans,rot}} = (x_i, y_i, z_i)^{\mathrm{T}}$ were transformed to spherical coordinates $\mathbf{r}_i^{\text{trans,rot}} = (r, \theta, \phi)^{\mathrm{T}}$.

Third, the rotation matrix

$$\mathbf{R}(\phi) = \begin{bmatrix} 1 & 0 & 0 \\ 0 & 1 & 0 \\ 0 & 0 & 1 - \phi^{\text{ref2}} \end{bmatrix}, \tag{6}$$

rotates all position vectors $\mathbf{r}_i^{\text{trans,rot}}$ around the $z$-axis, such that $C_\alpha$(Gly10) lies in the $zx$-plane. $\phi^{\text{ref2}}$ is the azimuth angle of the position vector of $C_\alpha$(Gly10) after the coordination transform.

As consequence, any conformation of the tWW domain was represented by the following reaction coordinates:

- $(r, \theta, \phi)$: the position of $C_\alpha$(Tyr61) in spherical coordinates.
- $\alpha$: the torsion angle defined by $C_\alpha$(His17), $C_\alpha$(Tyr20), $C_\alpha$(Thr58), $C_\alpha$(Tyr61)
- $\gamma$: the angle defined by $C_\alpha$(Thr58), $C_\alpha$(Tyr61) and $C_\alpha$(His17).

###### 2.2.4 Common-nearest-neighbor clustering

The data set of compact structures of apo-h-FBP21 tWW ($31.51 \cdot 10^6$ structures) was clustered in the space of the reaction coordinates ($r, \theta$, $\phi$, $\gamma$, $\alpha$) using the implementation of our group for the common-nearest-neighbor clustering (CommonNN) algorithm.[21,22] This implementation, called *CommonNN clustering*, is part of the python module `scikit-learn-extra` and available from GitHub (source: `https://github.com/janjoswig/CommonNNClustering`, documentation: `https://scikit-learn-extra.readthedocs.io/en/stable/generated/sklearn_extra.cluster.CommonNNClustering.html`).

Considering chronologically every 1000th structure, hierarchical clustering was performed.[23] To circumvent circular coordinates, $\phi$ was transformed into sin($\phi$) and cos($\phi$), likewise for $\theta, \alpha, \gamma$. The distance $r$ was scaled to the interval [0, 1].

The CommonNN clustering algorithm[21,22] identifies clusters based on the number of shared neighbours between two data points where the neighbourhood is defined by the radius $R$. The minimum number of neighbours that are needed for two data points to be in the same cluster is set by the parameter $N$. Additionally, small clusters can be classified as noise by specifying a minimal cluster size $M$. The cluster parameters and the percentage of data points classified as noise for each round of clustering is reported in Table S3. Overall, the clustering yielded 45 clusters that cover 41% of the cluster data set. For each cluster, one representative structure was selected respectively. These structures show the smallest backbone RMSD to all other members of their cluster.

Table S3: Overview of the cluster parameters required for applying the CommonNN clustering algorithm on the MD data of the apo-h-FBP21 tWW.

| hierarchy | 0 | 1 | | | | 2 | | | | | 3 | | |
|---|---|---|---|---|---|---|---|---|---|---|---|---|---|
| $R$ | 0.3 | 0.3 | 0.3 | 0.25 | 0.25 | 0.3 | 0.25 | 0.25 | 0.25 | 0.21 | 0.25 | 0.25 | 0.21 |
| $N$ | 29 | 39 | 39 | 39 | 37 | 46 | 46 | 46 | 46 | 39 | 46 | 52 | 47 |
| $M$ | 23 | 32 | 11 | 32 | 32 | 32 | 32 | 32 | 32 | 32 | 32 | 32 | 32 |
| noise | 34% | 13% | – | 27% | 19% | 9% | 31% | 26% | 14% | 23% | 14% | 9% | 36% |

Please note, due to the fact that on each level (> 0) different children of the next lower level were clustered, it is possible that the same combination of $R$, $N$ and $M$ can yield different clustering results.

###### 2.2.5 Backbone RMSD between apo-h-FBP21 tWW CommonNN cluster structures

For the calculation of the RMSD between the CommonNN clusters of the apo-h-FBP21 tWW, GROMACS 2019 release version[16] was used. First, a reference structure was extracted from a cluster $j$ using the tool `gmx trjconv` its cluster trajectory. Then, after a least-square fit on the $C_\alpha$ atoms of the h-FBP21 tWW, the backbone $\text{RMSD}_{ij}$ was calculated with `gmx rms` for the cluster $i$ of interest using the extracted structure of cluster $j$ as reference structure ('-s' flag). We calculated the $\text{RMSD}_{ij}$ for each single structure of cluster $j$ and the trajectory of cluster $i$ respectively. The average RMSD between cluster $i$ and $j$ was then computed as follows:

$$\mu\left(\text{RMSD}_{ij}\right) = \frac{1}{N_i N_j} \sum_{h=1}^{N_i} \sum_{k=1}^{N_j} \text{RMSD}_{i_h j_k} \tag{7}$$

with $N_i$, $N_j$ as the amount of structures in clusters $i$ and $j$. The matrix containing all average $\text{RMSD}_{ij}$ values is visualised in Fig. S2.

###### 2.2.6 Hydrogen bond analyses

Hydrogen bond analyses were performed with GROMACS 2016.5 simulation package using `gmx hbond`. After assigning each atom to either the side or the main chain of its residue with `gmx make_ndx`, all interactions were classified as side or main chain interactions of the involved residues. In case of the tWW, the hydrogen bonds were further classified in interdomain, i.e. the involved residues do not belong to the same WW domain, and intradomain, both residues belong to one WW domain, hydrogen bonds. For the apo-h-FBP21 tWW, we observed 631 intradomain and 1063 interdomain hydrogen bonds.

All hydrogen bonds were evaluated on the basis of their relative occurrence over the amount of considered structures, e.g. all structures of a cluster or all structures of an entire simulation. Hydrogen bonds with a relative occurrence greater than 50% were considered significant. Fig. S3 shows the populations of the interdomain hydrogen bonds for the apo-h-FBP21 tWW that were highly populated (> 50%) in at least one cluster (152 out of 1063).

#### 2.3 Docking experiments with HADDOCK protocol

##### 2.3.1 Explanation of HADDOCK

For each docking experiment, HADDOCK2.2 provides a series of proposals for the complex structure, where each proposal consists of a bundle of four closely related structures. All proposals are ranked on the basis of the HADDOCK scoring function, a weighted sum of energy-like contributions:[24]

$$\mathrm{HS} = 1.0\, E_{\mathrm{vdW}} + 0.2\, E_{\mathrm{elec}} + 1.0\, E_{\mathrm{desolv}} + 0.1\, E_{\mathrm{AIR}}\,, \qquad (8)$$

where $E_{\mathrm{vdW}}$ represents the van der Waals interactions, $E_{\mathrm{elec}}$ represents the Coulomb interactions, $E_{\mathrm{desolv}}$ is an empirical desolvation term and $E_{\mathrm{AIR}}$ is a soft-square harmonic potential used to impose restraints (e.g. from experimental data).[24–26] As it combines energy terms with empirical terms, the included HADDOCK score is not a measure for the binding affinity,[25,27,28] but it can be used to deduce information about the probable stability of the complex. Due to the possible integration of experimental data or information about binding relevant residues, the accuracy of the HADDOCK protocol, however, is enhanced in comparison to non-integrative docking approaches.[27,28] We therefore used it as a tool to generate possible complex structures.

##### 2.3.2 Usage of HADDOCK

Using the HADDOCK2.2 webserver,[14] a structure of the SmB2 ligand taken from Ref. 29, was docked to the representative cluster structures of the apo-h-FBP21 tWW obtained with the CommonNN clustering algorithm.
The following residues were set as active:

- tWW: Val8, Glu9, Gly10, Tyr18, Tyr20, Ser27, Trp29, Val49, Glu50, Gly51, Tyr59, Tyr61, Ser68, Trp70
- SmB2: Met1, Gly2, Met3, Pro4, Pro5, Pro6, Gly7, Met8, Arg9, Pro10, Pro11, Pro12, Pro13, Gly14, Met15, Arg16, Gly17, Leu18, Leu19

where Tyr18, Ser27, Trp29, Tyr59, Ser68 and Trp70 are part of the XP grooves in the tWW, and Pro4, Pro5, Pro6, Arg9, Pro10, Pro11, Pro12, Pro13, Arg16 make up the proline-rich sequence in the SmB2 ligand. All remaining residues were defined as passive. For all 45 representative h-FBP21 tWW structures, the docking returned proposals for the h-FBP21 tWW / SmB2 complex. We saved the two proposals with the lowest HADDOCK scores for each tWW structure (i.e. 90 proposals). Since each HADDOCK proposal consists of a bundle of four h-FBP21 tWW / SmB2 complex structures, we obtained in total $90 \cdot 4 = 360$ h-FBP21 tWW / SmB2 complex structures.

#### 2.4 Analysis of the h-FBP21 tWW / SmB2 complex simulations

In preparation for the analyses, all trajectories were modified as described in 2.2.

##### 2.4.1 Hydrogen bond analyses

Hydrogen bonds were analysed as described in 2.2.6. Besides the intra- and interdomain hydrogen bonds, two further different types of hydrogen bonds were distinguished: intramolecular, *i.e.* within the h-FBP21 tWW, and intermolecular, between the SmB2 ligand (mutated, WT) and the h-FBP21 tWW.

For the simulated HADDOCK proposals (1-15), we observed 1830 hydrogen bonds, of which 653 are intermolecular, 472 are interdomain and 624 are intradomain hydrogen bonds. The populations of the significant hydrogen bonds (population $> 50\%$) are visualised in Fig. S10 (44 interdomain hydrogen bonds) and Fig. S6 (31 intermolecular hydrogen bonds).

For the complexes 6 and 14 (mutated, WT), we observed 1284 different hydrogen bonds, of which 316 are intermolecular, 362 are interdomain and 606 are intradomain hydrogen bonds. The significant hydrogen bonds (population $> 50\%$) are presented in our study (see results section 2.3).

##### 2.4.2 Distances between XP grooves and proline-rich motifs

To classify the orientation of the ligand relative to the tWW (parallel, antiparallel), four groups of $C_\alpha$ atoms were defined:

1) XP1: Tyr18, Ser27, Trp29

2) XP1: Tyr59, Ser68, Trp70

3) PRM1: Pro4, Pro5, Pro6

4) PRM2: Pro10, Pro11, Pro12, Pro13

where XP1 and XP2 represent the XP grooves in either WW domain 1 or 2 of the h-FBP21 tWW structure, and PRM1 and PRM2 represent the two proline-rich motifs in the SmB2 ligand (mutated, WT). For each group $A$, the geometric centers $\mathbf{r}_A^{\text{geo}}$ were calculated as follows:

$$\mathbf{r}_A^{\text{geo}} = \frac{1}{N_A} \sum_{i=1}^{N_A} \mathbf{r}_{i,A}, \tag{9}$$

where $N_A$ is the number of atoms of group $A$.

Between all four geometric centers, the Euclidean distance was calculated with:

$$d(A,B) = \mid \mathbf{r}_B^{\text{geo}} - \mathbf{r}_A^{\text{geo}} \mid . \tag{10}$$

A complex is classified ‘parallel’, if the Euclidean distances $d$(XP1,PRM1) and $d$(XP2,PRM2) are smaller than $d$(XP1,PRM2) and $d$(XP2,PRM1), and ‘antiparallel’ *vice versa.*

##### 2.4.3 Extraction of representative complex structures

Based on the Euclidean distances between the geometric centers, complex structures were extracted if:

$$|d(\text{XP1}, \text{PRM}) - d(\text{XP2}, \text{PRM})| = 0, \tag{11}$$

with $d(\text{XP1/2}, \text{PRM})$ as the distance between the geometric center of the group XP1 or XP2 and the geometric center of the respective closest proline-rich motif.

#### 2.5 Protein preparation and construct cloning

The construct used for the h-FBP21 tWW was described in previous studies.[29] The h-FBP21 single domain WW1 construct was obtained by inserting residues 123-159 of h-FBP21 in petM11 vector, using NcoI/XholI as restriction sites. The h-FBP21 single domain WW2 construct was achieved by insertion of residues 158-200 in pGEX-4T-1 via BamHI/EcoRI. The construct h-FBP21 tWW-R6A with the substitution of Arginine 127 with Alanine was achieved by Quick Change Mutation. Escherichia Coli BL21(DE3) were transformed with the plasmids and expanded in M9 minimal medium containing 750 mg/l of $^{15}$N enriched

Ammonium Chloride and, in order to acquire 3D spectra, also 2 g/l of $^{13}$C D-glucose. For h-FBP21 tWW-R6A a site selectively labeling strategy was performed to implement the protein assignment. Protein expression was induced with IPTG at OD≈0.6 and incubated overnight at 18 °C. Protein purification was performed via affinity chromatography using HisTrap HP or GSTrap FF (GE Healthcare) based on the tag expressed in the N-terminal part of the protein, which was subsequently removed via Thrombin or TEV digestion. Further purification was achieved with size exclusion chromatography (Superdex 75 10/300 GL) using phosphate buffer (150 mM NaCl, 50 mM $KH_2PO_4$, 1 mM EDTA at pH 6.7).

#### 2.6 NMR spectroscopy

NMR measurements were performed on a Bruker Avance III 700 MHz spectrometer equipped with a 5 mm triple-resonance cryoprobe. To process the spectra, Topspin 3.2 software (Bruker) was used, whereas the spectra evaluation was performed with CcpNmr Analysis (version 2.4.2).[30] $^1$H-$^{15}$N-HSQC spectra were measured at 298 K with a protein concentration of 100 $\mu$M in phosphate buffer with 10% $D_2O$ v/v at pH 6.7 with 8 scans and 1024 ($^1$H) and 128 ($^{15}$N) data points. To investigate structural and conformational changes between the h-FBP21 tWW and the single WW domain 1 and WW domain 2 (see results section 2.1) as well as between the h-FBP21 tWW in apo- and holo-state (see Fig. S8), $^1$H-$^{15}$N-HSQC spectra were compared and the $^1$H-$^{15}$N chemical shift differences were calculated as follows:

$$\Delta\delta = \sqrt{(\Delta^1\mathrm{H})^2 + (0.15 \cdot \Delta^{15}\mathrm{N})^2}, \qquad (12)$$

with $\Delta^1$H and $\Delta^{15}$N as the differences in the $^1$H- and $^{15}$N-signal respectively. Shifts bigger than the average plus standard deviation ($\mu(\Delta\delta) + \sigma(\Delta\delta)$) were considered significant. The backbone assignment for the singular WW domains of the h-FBP21 tWW was obtained from standard triple resonance spectra. HNCA spectra of WW domain 1 and WW domain 2 were acquired with 16 scans and 1024x96x96 and 1028x88x96 complex data points ($^1$H,$^{13}$C,$^{15}$N) respectively. For the WW domain 2, an HN(CO)CA spectrum was additionally measured with 24 scans 1024x88x128 complex data points and 60% non-uniform sampling to reduce measurement time. For the h-FBP21 tWW in complex with the SmB2 ligand, a $^{1}$H-$^{15}$N-NOESY-HSQC spectrum was acquired with 32 scans, 1024x72x120 data points and 80 ms mixing time on a Bruker Avance 750MHz spectrometer equipped with a triple-resonance cryo-probe.

#### 2.7 Isothermal Titration Calorimetry

The Isothermal Titration Calorimetry measurements were performed at 298.15 K with the GE device MicroCaliTC200. The cell was filled up with 100 $\mu$M of protein, both h-FBP21 tWW and its mutant tWW-R6A were previously dialysed in phosphate buffer. The titration was achieved with 76 injections of the proline-rich ligand of the core-splicing protein SmB/B' (1.3 mM, Ac-GTPMGMPPPGMR-PPPPGMRGLL-NH2). The dissociation constant values ($K_{\mathrm{D}}$) were obtained by plotting the integrated area peaks against the molar ratio, the data was fitted using "One Set of Sites". Thermodynamic parameters differ from previous publication[29] due to different condition used.

### Present addresses

⊤D.G.: Institute of Chemistry and Biochemistry, Laboratory of Structural Biochemistry, Freie Universität Berlin, 14195 Berlin, Germany
